# Cyfip2 controls the acoustic startle threshold through FMRP, actin polymerization, and GABA_B_ receptor function

**DOI:** 10.1101/2023.12.22.573054

**Authors:** Jacob C. Deslauriers, Rohit P. Ghotkar, Lindsey A. Russ, Jordan A. Jarman, Rubia M. Martin, Rachel G. Tippett, Sureni H. Sumathipala, Derek F. Burton, D. Chris Cole, Kurt C. Marsden

**Affiliations:** Department of Biological Sciences, North Carolina State University, Raleigh, North Carolina, USA; Center for Human Health and the Environment (CHHE), North Carolina State University, Raleigh, North Carolina, USA; Putnam Associates, Boston, Massachusetts, USA; Department of Pharmacology & Physiology, Georgetown University, Washington D.C., USA; Department of Physiology and Biophysics, Boston University, Boston, MA, USA; U.S. Environmental Protection Agency, Raleigh-Durham-Chapel Hill, North Carolina, USA; Biogen, Durham, North Carolina, USA

## Abstract

Animals process a constant stream of sensory input, and to survive they must detect and respond to dangerous stimuli while ignoring innocuous or irrelevant ones. Behavioral responses are elicited when certain properties of a stimulus such as its intensity or size reach a critical value, and such behavioral thresholds can be a simple and effective mechanism to filter sensory information. For example, the acoustic startle response is a conserved and stereotyped defensive behavior induced by sudden loud sounds, but dysregulation of the threshold to initiate this behavior can result in startle hypersensitivity that is associated with sensory processing disorders including schizophrenia and autism. Through a previous forward genetic screen for regulators of the startle threshold a nonsense mutation in *Cytoplasmic Fragile X Messenger Ribonucleoprotein (FMRP)-interacting protein 2* (*cyfip2*) was found that causes startle hypersensitivity in zebrafish larvae, but the molecular mechanisms by which Cyfip2 establishes the acoustic startle threshold are unknown. Here we used conditional transgenic rescue and CRISPR/Cas9 to determine that Cyfip2 acts though both Rac1 and FMRP pathways, but not the closely related FXR1 or FXR2, to establish the acoustic startle threshold during early neurodevelopment. To identify proteins and pathways that may be downstream effectors of Rac1 and FMRP, we performed a candidate-based drug screen that indicated that Cyfip2 can also act acutely to maintain the startle threshold branched actin polymerization and N-methyl D-aspartate receptors (NMDARs). To complement this approach, we used unbiased discovery proteomics to determine that loss of Cyfip2 alters cytoskeletal and extracellular matrix components while also disrupting oxidative phosphorylation and GABA receptor signaling. Finally, we functionally validated our proteomics findings by showing that activating GABA_B_ receptors, which like NMDARs are also FMRP targets, restores normal startle sensitivity in *cyfip2* mutants. Together, these data reveal multiple mechanisms by which Cyfip2 regulates excitatory/inhibitory balance in the startle circuit to control the processing of acoustic information.

## Introduction

To navigate their environments to find food and avoid predation, animals must be able to filter out extraneous stimuli but respond appropriately to salient ones, a process known as sensorimotor gating. Specific attributes of a stimulus can trigger a response; for visual stimuli the luminance, size, and speed of the stimulus determine if escape and reorientation responses are initiated [1–3]. Similarly, the intensity and frequency of acoustic stimuli determine whether a response is made [4,5]. One way in which animals can control their responses to sensory stimuli is by establishing a behavioral threshold such that when one or more of these stimulus attributes reaches a critical value a specific behavioral response is initiated. Behavioral thresholds are a fundamental mechanism of sensorimotor gating used across the animal kingdom to regulate a wide range of behavioral responses including both collective responses, such as fanning behaviors for hive climate regulation in bees [6,7] and shoaling behavior in fish [8,9], as well as individual responses to odors [10–13], tactile stimuli [14–17], changes in luminance and contrast of visual stimuli [1,18–21], and sound frequency and intensity in mammals and fish [5,22–24]. That behavioral over-responsiveness to visual, tactile, and acoustic stimuli is observed across a number of neuropsychiatric conditions including autism, anxiety, and schizophrenia [4,22,24–28] highlights the importance of setting such behavioral thresholds at an appropriate level. Yet our knowledge of the molecular mechanisms that both establish and maintain behavioral thresholds is limited.

Previously, to identify genes that regulate the threshold for initiating the acoustic startle response, a highly conserved behavior initiated following sudden loud sounds that may indicate danger [5,29–31], we conducted a standard 3-generation, ENU-based forward genetic screen in larval zebrafish [32]. We identified a set of five mutant lines that display acoustic startle hypersensitivity, and through whole-genome sequencing of the *triggerhappy* mutant line, we identified a causal, nonsense mutation in *cytoplasmic Fragile X Messenger Ribonucleoprotein (FMRP)-Interacting protein 2* (*cyfip2).* Cyfip2 was first identified as an interactor of FMRP and the elongation initiation factor 4E (eIF4E), through which it participates in translational repression of many target transcripts [33,34]. Cyfip2, but not the closely related Cyfip1, can also bind the Fragile X-related proteins FXR1 and FXR2, but the function of these interactions is unknown [33]. Additionally, Cyfip2 interacts with the activated form of the small Rho GTPase Rac1, and it is a member of the WAVE Regulatory Complex (WRC) in which it helps regulate Arp2/3 activation and branched actin polymerization [35–41]. Cyfip2 is vital for proper neuronal migration and cell movement, axonal growth and guidance, as well as synapse formation and function in flies, mice, and zebrafish [32,33,37,42–47]. Homozygous *cyfip2* mutations are embryonically lethal in mammals and fatal after 7-8 days post-fertilization (dpf) in zebrafish [32,42,44,48]. Despite its key role in multiple aspects of neurodevelopment, the links between how Cyfip2 regulates RNA translation, actin polymerization, and behavior have not been defined. Our previous work demonstrated that loss of Cyfip2 causes acoustic startle hypersensitivity that is reversible upon transgenic expression of GFP-tagged Cyfip2, alters the morphology but not the electrophysiological properties of the startle command-like Mauthner cells (M-cells), and causes hyperexcitability of the spiral fiber neurons (SFNs), a set of hindbrain excitatory interneurons that project to the M-cell axon hillock [32]. It is unclear, however, whether Cyfip2 acts via Rac1-mediated actin polymerization or through FMRP-mediated translational repression to control the startle threshold. Furthermore, the downstream molecular changes that directly modulate the function of the startle circuit have not been identified.

In this study we used an inducible rescue approach in *cyfip2* mutant zebrafish larvae to demonstrate that both Cyfip2’s Rac1 and FMRP interactions are required for establishing the acoustic startle threshold during early neurodevelopment. Using CRISPR-Cas9 gene knockdown we find that FXR1 and FXR2 are dispensable for startle regulation and that Cyfip2 acts through FMRP. Furthermore, with a candidate-based pharmacological approach we show that Cyfip2 mediates Arp2/3-induced branched actin polymerization and may modulate N-methyl-D-aspartate receptors (NMDARs) to alter neuronal function in the acoustic startle circuit. Finally, we performed discovery proteomics to define molecular pathways disrupted by loss of Cyfip2 *in vivo*. Our results indicate roles for Cyfip2 in mitochondrial function, oxidative phosphorylation, and inhibitory Gamma-Aminobutyric Acid (GABA) receptor signaling. We confirmed the functional importance of this last finding using the GABA_B_ receptor agonist baclofen, which rescues *cyfip2* mutants’ hypersensitivity. Together these data establish a novel pathway that links Cyfip2, actin dynamics, RNA translation, and excitatory/inhibitory balance in the control of acoustic responsiveness.

## Results

### Cyfip2 establishes the acoustic startle threshold through both Rac1 and FMRP during early neurodevelopment

Cyfip2 has four known protein-interaction domains [49] (Fig. 1A), and it can act through Rac1 to promote actin polymerization (Fig. 1B) and FMRP to regulate RNA translation (Fig. 1C). *cyfip2^p400^* mutants have a single base pair transversion (nt1024; T to A) resulting in a premature stop codon at amino acid position 343 (Fig. 1A) [32]. *cyfip2^p400^* mutant zebrafish larvae (5 dpf) were previously shown to display acoustic startle hypersensitivity that could be rescued by expressing Cyfip2 at 30 hours post fertilization (hpf) using a stable heatshock-inducible transgenic line, *Tg(hsp70:cyfip2-EGFP)* [32]. We replicate those findings here, using a 60-stimulus assay consisting of 10 trials at each of 6 intensities with a 20 second (s) interstimulus interval (ISI) to measure startle sensitivity (Fig. 1D). A 40-min heatshock at 30 hpf restores normal sensitivity in transgenic (*Tg*+) but not in non-transgenic (*Tg*-) *cyfip2* mutants (Fig. 1D,E). Previous studies have established that C179R and K723E amino acid substitutions prevent Cyfip2 from binding with Rac1 and FMRP, respectively [34,39,45]. To determine if Cyfip2 engages Rac1-mediated actin regulation and/or FMRP/eIF4E-mediated translational repression pathways to establish the acoustic startle threshold, we induced C179R (Δ*Rac1*) and K723E (Δ*FMRP*) point mutations in the *Tg(hsp70:cyfip2-EGFP)* construct and created stable transgenic lines for each (Fig. 1A). We expressed either wildtype or mutant (Δ*Rac1*; Δ*FMRP)* versions of Cyfip2 in mutants at 30 hpf with a 40-minute heatshock at 38°C, followed by acoustic startle testing at 5 dpf. While expression of wildtype *Tg(hsp70:cyfip2-EGFP)* at 30 hpf rescues mutant hypersensitivity, expression of Δ*Rac1* or Δ*FMRP* versions of Cyfip2 in mutants was insufficient to rescue mutant hypersensitivity (Fig. 1E).

**Figure 1.**
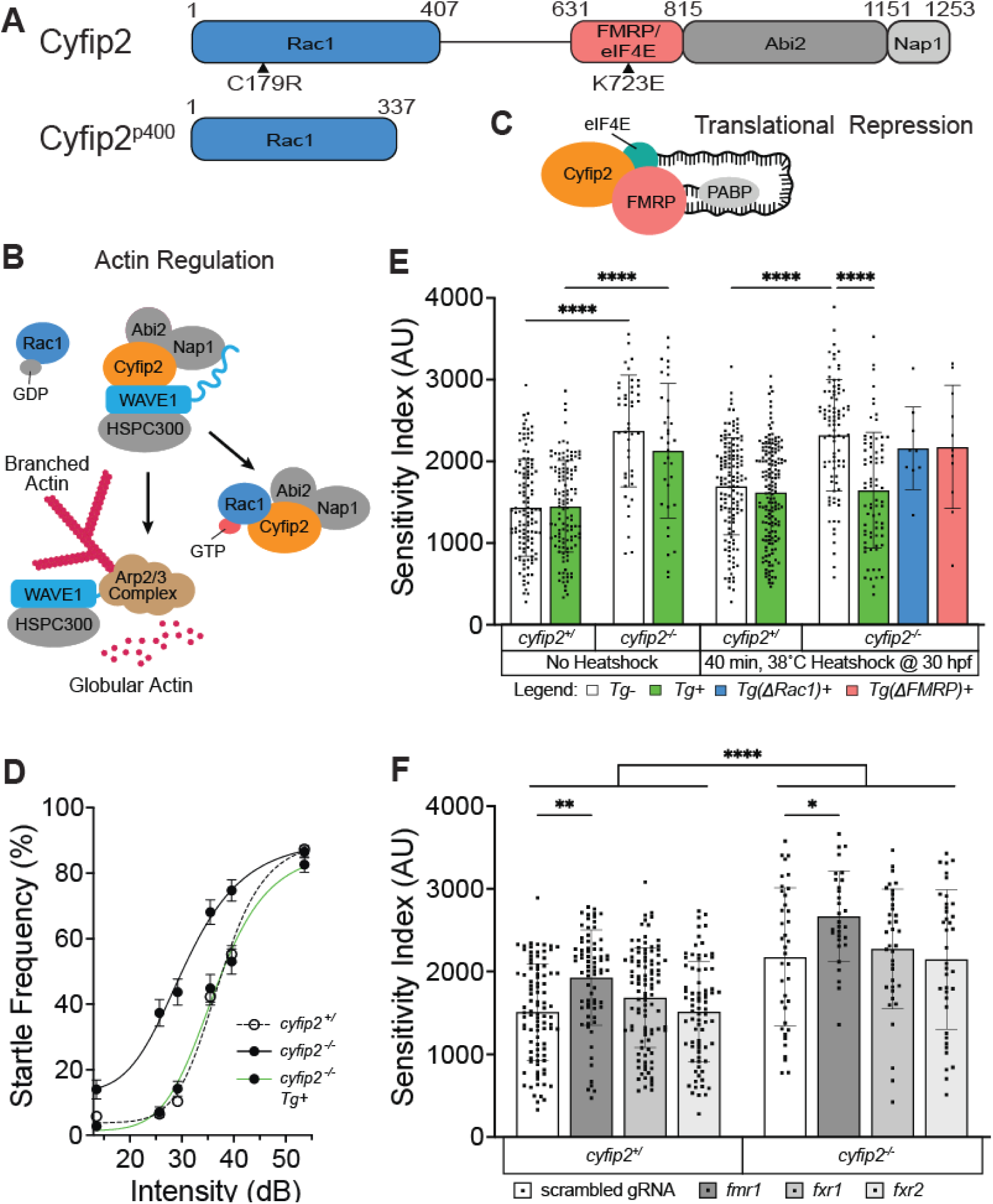
Cyfip2 establishes the acoustic startle threshold through Rac1 and FMRP. (A) Cyfip2 protein interacting domain diagram of wildtype (top) and mutant (bottom) Cyfip2 proteins. Black arrowheads indicate the positions of induced mutations in Cyfip2, eliminating the Rac1-(C179R) or FMRP/eIF4E (K723E)-binding capacity of Cyfip2. (B) Cyfip2 actin regulatory pathway wherein Cyfip2 (orange) upon stimulation by Rac1-GTP triggers WAVE1 activation, Arp2/3-complex initiation and branched actin nucleation. (C) Cyfip2 translational repression pathway in which Cyfip2, eIF4E (teal), and FMRP (pink) along with the poly-A binding protein (PABP; gray), sequester neurodevelopmentally important mRNAs from being translated. (D) Average startle frequency (%) after 10 trials at 13.6, 25.7, 29.2, 35.5, 39.6 and 53.6 dB for 5 dpf *cyfip2* siblings (+/) and mutant (−/−) larvae heatshocked at 30 hpf for 40 minutes at 38°C. The average startle frequency curve for cyfip2 siblings (+/; open circles, dashed line), cyfip2 mutants (−/−; closed circles, solid line) and cyfip2 mutants harboring the *Tg(hsp70:cyfip2-EGFP)*+ transgene (−/−; Tg+; closed circles, solid green line). (E) Sensitivity indices, calculated as the area under the startle frequency curves, for 5 dpf cyfip2 siblings and mutants, following a 40-minute heatshock at 30 hpf to express either wildtype (Tg+; green), Rac1-(ΔRac1+; blue) or FMRP/eIF4E-(ΔFMRP+; pink) binding deficient versions of Cyfip2-EGFP. Comparisons were made to both non-transgenic (Tg-) and non-heatshocked controls. All indices (mean ± SD) compared using a Kruskal-Wallis test with Dunn’s multiple comparisons correction; p**** < 0.0001. (F) Sensitivity indices for 5 dpf *cyfip2* sibling (+/) and mutant (−/−) larvae following 1-cell stage injection with CRISPR-Cas9 and a single, scrambled guide RNA (gRNA) or dual gRNA cocktails targeting *fmr1*, *fxr1*, or *fxr2*. scrambled gRNA injected (white bar, closed circles); fmr1 gRNA injected (dark gray bar closed circles); *fxr1* gRNA injected (medium gray bar; closed circles); *fxr2* gRNA injected (light gray bar, closed circles). Comparisons were made both within genotype and between genotypes by condition. All indices (mean ± SD) compared using an Ordinary one-way ANOVA with Sidak’s multiple comparisons correction; p* < 0.05; p** < 0.01; p**** < 0.0001.

*cyfip2* mutants also display a number of kinematic defects in their performance of the startle response [32]. To determine if Rac1 and FMRP binding are also required for Cyfip2 to regulate startle kinematics, we analyzed startle latency, duration, head turn angle (C1 angle), and total distance traveled during the response in *Tg*- and *Tg*+ fish after the same 40-min heatshock protocol at 30 hpf (Fig. S1). Heatshock induction of wildtype Cyfip2 restored normal latency, duration, and C1 angle, but not distance traveled (Fig. S1). All kinematic defects remained in *cyfip2* mutant larvae expressing Δ*Rac1*-Cyfip2 (Fig. S1), but expression of Δ*FMRP*-Cyfip2 was sufficient to rescue startle duration and induced a trend toward rescue of startle latency and C1 turn angle (Fig. S1). These data suggest that Rac1 binding is required for all aspects of Cyfip2-mediated startle regulation, while FMRP binding is required to regulate the startle threshold but is largely dispensable for startle kinematics.

One possible explanation for these results is that expression levels may differ between the three heatshock transgenic lines. We therefore measured expression levels of each transgenic Cyfip2-GFP protein 6 hours after a 40-min heatshock by fluorescence intensity (Fig. S2A). The Δ*Rac1* and Δ*FMRP* lines displayed GFP expression that was not significantly different than the wildtype *Tg+* line but which trended lower. To induce expression of wildtype Cyfip2-GFP at levels more comparable to the Δ*Rac1* and Δ*FMRP* lines after 40-minute heatshock, we delivered a 15-min heatshock at 30 hpf in the wildtype *Tg+* line. The 15-min heatshock reduced peak Cyfip2-GFP expression to levels comparable to or below that of the Δ*Rac1* and Δ*FMRP* lines, and this level of expression was also sufficient to rescue acoustic startle sensitivity in *cyfip2* mutants (Fig. S2A-C). Thus, the level of transgene expression cannot account for the failure of the Δ*Rac1* and Δ*FMRP* constructs to rescue startle phenotypes, and these findings support our conclusion that Cyfip2 utilizes both Rac1- and FMRP-mediated pathways to establish the acoustic startle threshold.

### Cyfip2 acts through FMRP but not FXR1/2 to establish the acoustic startle threshold

We previously found that FMRP is not required to establish a normal startle threshold, as *fmr1^hu2787^* mutants’ startle sensitivity is unaffected [32]. Having established that the K723 residue that Cyfip2 uses to bind FMRP is required for normal startle sensitivity, however, we hypothesized that Cyfip2 may instead interact with other members of the Fragile X protein family, Fragile X-related proteins 1 and 2 (FXR1/2), to establish the acoustic startle threshold [33]. We designed pairs of CRISPR guide RNAs (gRNAs) targeting each of the *fmr1, fxr1* and *fxr2* genes. We injected *fmr1-*, *fxr1-* or *fxr2*-specific CRISPR-Cas9 cocktails into 1-cell stage *cyfip2* sibling and mutant embryos and measured acoustic startle sensitivity in these crispant larvae at 5 dpf (Fig. 1F). Highly efficient CRISPR-induced mutagenesis was observed for all 3 genes, with 4 of the 6 gRNAs inducing edits as confirmed by PCR amplification and Sanger sequencing (Fig. S3A-F). Quantitative PCR confirmed that mRNA expression of all three genes was strongly reduced by CRISPR/Cas9 injection (Fig. S3G). FMRP crispants had significantly increased startle sensitivity compared to larvae injected with a scrambled gRNA plus Cas9, and sensitivity of FMRP crispants was even further increased in the *cyfip2* mutant background (Fig. 1F). However, startle sensitivity was unaltered in FXR1 or FXR2 crispants in either *cyfip2* siblings or mutants (Fig. 1F). These data indicate that FMRP, but not FXR1/2, regulates the startle threshold. The discrepancy between our hypersensitive *fmr1* crispants and prior analysis of non-hypersensitive *fmr1^hu2787^* mutants could be due to genomic adaptation that has been reported in the *fmr1^hu2787^*mutant line that may partially compensate for the loss of FMRP [50]. In support of this, an independently created CRISPR-generated *fmr1* mutant line displayed additional behavioral and developmental phenotypes not originally seen in the *fmr1^hu2787^* line [51]. *fmr1^hu2787^* mutant larvae have been found to express other autism-related phenotypes, however, such as altered social behavior and preference for reduced sensory stimulation [52], as well as increased brain activity in response to acoustic stimulation [53]. Together our conditional rescue and *fmr1* crispant data provide clear evidence that Cyfip2 acts in part through FMRP to establish the acoustic startle threshold.

### Cyfip2 can acutely maintain the acoustic startle threshold through Rac1 and FMRP pathways

Our previous study found that Cyfip2 is important for both establishing the acoustic startle threshold during early neurodevelopment, and for actively modulating the threshold later in development between 4 and 6 dpf [32]. Here we sought to define a critical window for Cyfip2 expression in regulating the startle threshold using our heatshock transgenic lines. Our results show that while a 40-minute heatshock to induce expression of wildtype Cyfip2 (*Tg+*) at 30 hpf is sufficient for behavioral rescue in 5 dpf *cyfip2* mutant larvae (Fig. 1D,E), a 40-min heatshock at 2, 3, or 4 dpf fails to rescue *cyfip2* mutant hypersensitivity (Fig. S4). Heatshock-induced expression at 5 dpf, 4 hours prior to testing, resulted in a trend toward rescue and a bi-modal distribution with some *cyfip2* mutants remaining hypersensitive and a second population showing restoration of normal sensitivity (Fig. 2A). These results are similar to our prior findings [32], and they suggest that Cyfip2 can not only function during the development of the startle circuit but can also actively maintain the circuit’s threshold after it has formed. Next, to determine if Cyfip2 employs both of its canonical pathways in maintaining the startle threshold after 4 dpf, we expressed either Δ*Rac1* or Δ*FMRP* versions of Cyfip2 in *cyfip2* mutants at 5 dpf with a 40-min heatshock followed by acoustic startle behavior testing 6 hours later. Similar to what we observed with the developmental heatshock (Fig. 1E), neither expression of the Δ*Rac1* nor the ΔFMRP version of Cyfip2 at 5 dpf rescued acoustic hypersensitivity of *cyfip2* mutants (Fig 2A). These findings suggest that Cyfip2 uses both its Rac1- and FMRP-mediated pathways to both establish and dynamically maintain the acoustic startle threshold throughout neurodevelopment.

**Figure 2.**
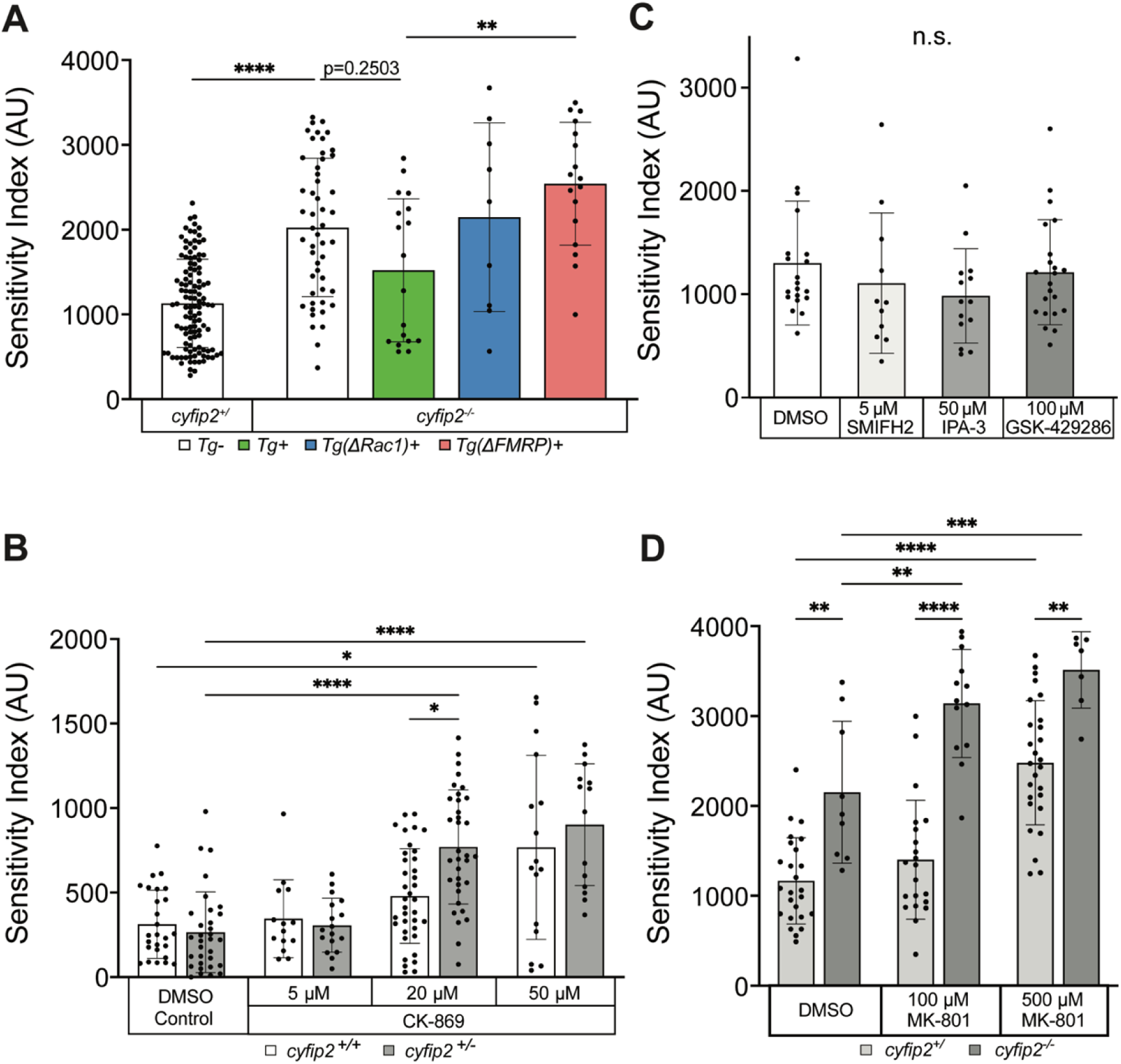
Cyfip2 acutely regulates branched actin polymerization and NMDARs to establish the acoustic startle threshold. (A) Sensitivity indices for 5 dpf *cyfip2* sibling (+/) and mutant (−/−) larvae, following a 40-minute heatshock at 120 hpf (5 dpf) to express either wildtype (Tg+; green), Rac1-(ΔRac1+; blue) or FMRP/eIF4E-(ΔFMRP+; pink) binding deficient versions of Cyfip2-EGFP. Comparisons were made to non-transgenic (Tg-), heatshocked sibling (+/) and mutant (−/−) controls. All indices (mean ± SD) compared using a Kruskal-Wallis test with Dunn’s multiple comparisons correction; p-values listed; p** < 0.01, p**** < 0.0001. (B) Sensitivity indices for 5 dpf *cyfip2* wildtype (+/+; white bar) and heterozygous (+/-; gray bar) larvae, treated for 30 minutes on d5 with 5, 20 or 50 µM CK-869. Comparisons were made both within genotype and within condition. All indices (mean ± SD) compared using a Kruskal-Wallis test with Dunn’s multiple comparisons correction; p* < 0.05; p**** < 0.0001. (C) Sensitivity indices for 5 dpf Tüpfel longfin (TL) larvae treated for 30 minutes on d5 with the highest, non-lethal doses the formin antagonist (SMIFH2; 5 µM), PAK3 antagonist (IPA-3; 50 µM) and ROCK antagonist (GSK429286; 100 µM). Comparisons were made between respective treatments and the DMSO controls. All indices (mean ± SD) were compared using a Kruskal-Wallis test with Dunn’s multiple comparisons correction; All comparisons made were non-significant (n.s.). (D) Sensitivity indices for 5 dpf *cyfip2* sibling (+/) and mutant (−/−) larvae, treated for 30 minutes on d5 with 100 or 500 µM MK-801. Comparisons were made both between genotypes within condition and between conditions by genotype. All indices (mean ± SD) were compared using an Ordinary one-way ANOVA with Tukey’s multiple comparisons correction. p** < 0.01; p*** < 0.001; p**** < 0.0001.

### Cyfip2 maintains the acoustic startle threshold through branched actin polymerization

Regulation of the actin cytoskeleton is a vital cellular process that within the context of the nervous system is critical for cell migration and movement, synapse formation, function and plasticity, receptor anchoring and trafficking, as well as axon growth and guidance [54]. Given our findings that Cyfip2 can act through the Rac1-WAVE pathway to establish and modulate the acoustic startle threshold, we hypothesized that Cyfip2-mediated branched actin polymerization specifically modulates the startle threshold. To test this hypothesis, we incubated 5 dpf *cyfip2* heterozygous and wildtype larvae in the Arp2/3 antagonist CK-869 at 5, 20, or 50 µM for 30 minutes, followed by acoustic startle testing. In control conditions, *cyfip2* heterozygotes display startle sensitivity equal to that of wildtypes (Fig. 2B), and incubation at 5 µM did not significantly alter startle sensitivity of either heterozygotes or wildtypes (Fig 2B). At 20 µM, wildtype larvae are unaffected by CK-869, but *cyfip2* heterozygotes have significantly increased startle sensitivity compared to controls (Fig 2B). 50 µM CK-869 caused both *cyfip2* wildtype and heterozygous larvae to become significantly hypersensitive compared to controls, phenocopying *cyfip2* mutants. Thus, Arp2/3-mediated, branched actin polymerization is necessary for acutely maintaining the acoustic startle threshold. That *cyfip2* heterozygotes display hypersensitivity at 20 µM but wildtypes do not indicates that a single functional copy of *cyfip2* is insufficient to maintain normal startle circuit function when branched actin polymerization is limited by a moderate dose of CK-869. We found similar results when exposing larvae to another Arp2/3 antagonist, CK-666, with both *cyfip2* heterozygotes and wildtypes phenocopying mutant hypersensitivity at a 50 µM concentration (Table S2). These findings support the conclusion that Cyfip2-dependent Arp2/3-mediated branched actin polymerization is necessary to maintain the acoustic startle threshold.

The actin cytoskeleton is dynamic and requires the action of both branched and unbranched actin regulatory pathways to maintain cellular structure and function. Unbranched filamentous (F) actin is polymerized from globular actin monomers by dimeric complexes of formin proteins, which bind at the barbed ends of new filaments and promote their elongation [55–58]. Another form of actin regulation involves the action of cofilin, an actin severing protein that cleaves existing filaments to create new barbed ends and increase the rate of actin turnover within the cell [59]. To determine whether unbranched actin and severing pathways play a role in regulating the acoustic startle threshold we incubated 5 dpf wildtype larvae in a formin antagonist, SMIFH2, or the cofilin disinhibitors, IPA-3 and GSK429286, for 30 minutes followed by acoustic startle testing. Treatment with 5 µM SMIFH2 or with 50 µM IPA-3 or 100 µM GSK429286 did not significantly affect startle sensitivity in wildtype larvae (Fig 2C). We also tested these drugs at higher concentrations, which were lethal after a 30-minute exposure, as well as longer exposures at lower concentrations, which had no effect on startle sensitivity (Table S1). Our data with the formin inhibitor SMIFH2 are in contrast to a recent finding showing that Formin 2B morpholino knockdown caused a decrease in Mauthner cell-mediated fast startle responses in zebrafish larvae [60]. That we did not observe any change in startle frequency could be due to the acute nature of our pharmacological approach as opposed to the morpholino-mediated developmental knockdown of Formin 2B. Our findings suggest that acute perturbations to unbranched actin filaments and actin turnover do not play a significant role in regulating the acoustic startle threshold. Altogether, these data further support our conclusion that Cyfip2-mediated, branched actin polymerization is a key pathway for acutely maintaining the acoustic startle threshold.

### Cyfip2 may regulate NMDA receptors to modulate the acoustic startle threshold

While we have established that Cyfip2 mediates the establishment and maintenance of the acoustic startle threshold through both branched actin and FMRP regulatory pathways, it is unclear what molecular mechanisms directly modulate the excitability of the startle circuit. To identify molecules that may be downstream effectors of Cyfip2 in modulating activity of the startle circuit, we conducted a candidate-based small-molecule screen with compounds previously shown to alter startle sensitivity in wildtype zebrafish larvae (Table 1) [61]. In this screen we incubated *cyfip2* wildtype, heterozygous, and mutant larvae in each compound for 30 min prior to and during acoustic startle testing. Consistent with previous findings, *N-* phenylanthrinilic acid (NPAA; Cl^-^ channel antagonist), Meclofenamic acid (MA; K^+^ channel and gap junction antagonist), Phenoxybenzamine (POBA; alpha-adrenergic receptor and calmodulin antagonist), Etazolate (ETAZ; phosphodiesterase 4 (PDE4) inhibitor), and MK-801 (N-methyl-D-aspartate receptor (NMDAR) antagonist) all increased startle sensitivity in a dose-dependent manner (Table 1). BMS204352, a different K^+^ channel antagonist, did not alter acoustic startle sensitivity at either 10 or 50 µM concentrations, and NSC-23766, a Rac1 antagonist, reduced sensitivity in siblings at 100 µM, but not *cyfip2* mutants.

**Table 1.**
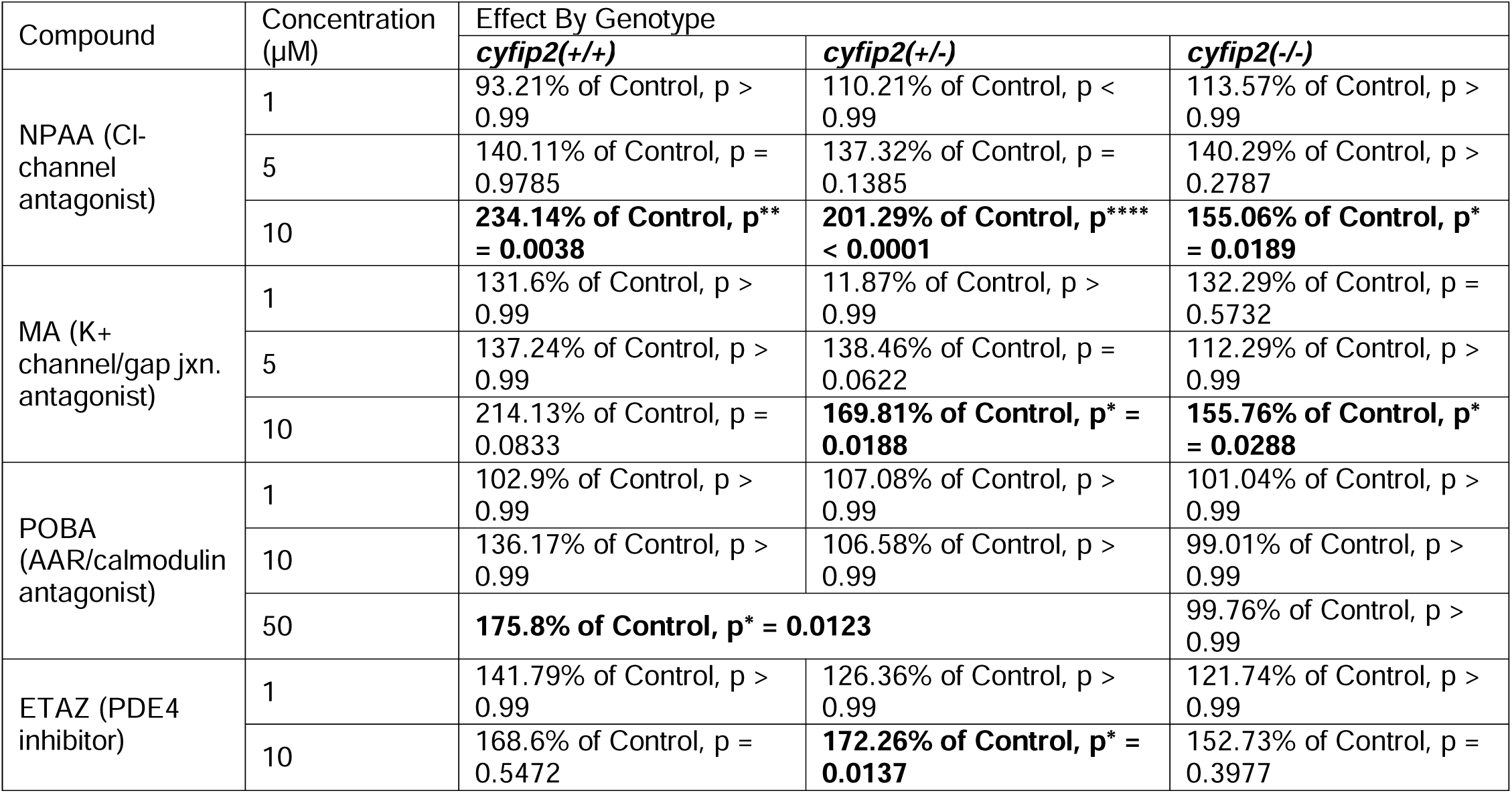
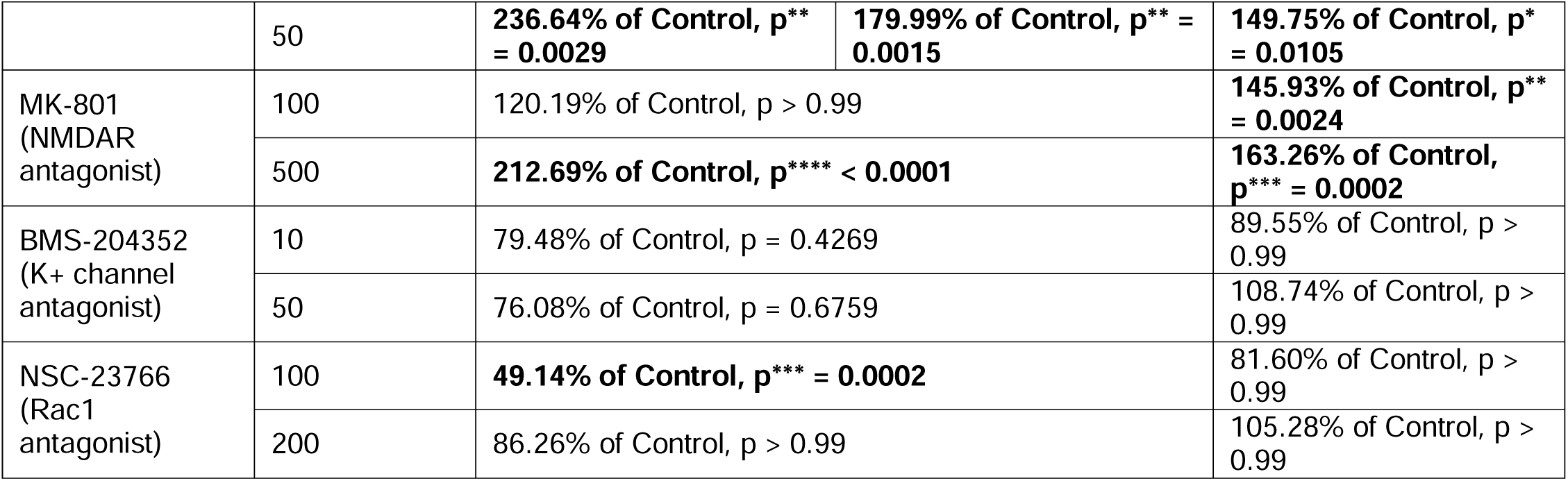
Cyfip2 may regulate NMDARs to control acoustic startle sensitivity. Mean startle index comparisons, listed as percentage (%) of the mean startle index of vehicle-treated controls by *cyfip2* genotype and drug concentration, for larvae treated with compounds targeting the indicated pathways [61] to increase acoustic startle sensitivity. All significant differences (p < 0.05) are listed (**bold**) for comparisons using a Kruskal-Wallis test and Dunn’s multiple comparisons correction. NPAA (*N*-phenylanthranilic acid); MA (meclofenamic acid); POBA (phenoxybenzamine); ETAZ (etazolate).

To determine whether any of the targeted pathways may be downstream of Cyfip2, we looked for conditions in which there was a clear *cyfip2* genotype-specific effect on sensitivity. The NMDA receptor blocker MK-801 showed the clearest such effect, with a low dose (100 µM) elevating startle sensitivity only in *cyfip2* mutants but not siblings (Table 1). Therefore, Cyfip2-mediated cytoskeletal and/or translational regulation may impact the expression and/or function of NMDA receptors within the startle circuit to modulate the acoustic startle threshold.

### Proteomic analysis reveals that Cyfip2-mediated regulation of GABA_B_ receptors is critical for startle sensitivity

To complement our candidate drug screen with an unbiased approach to identify proteins and molecular pathways regulated by Cyfip2, likely through its role in translational regulation, we conducted a proteomic analysis of *cyfip2* wildtype, heterozygous, and mutant larvae at 5 dpf. All larvae used were siblings and were genotyped by PCR and Sanger sequencing and then pooled in groups of 30 per genotype and snap frozen with liquid nitrogen. Three independent pools of 30 larvae were analyzed for each genotype. Protein lysates were submitted to the Molecular Education, Technology and Research Innovation Center (METRIC) at NC State University for protein digestion and LC-MS. Raw LC-MS files were processed and quantified using MaxQuant (Max Planck Institute of Biochemistry) and imported into Perseus software for transformation and identification of Differentially Expressed Proteins (DEPs) for subsequent Ingenuity Pathway Analysis (IPA).

Comparative analysis of *cyfip2* heterozygous and mutant versus wildtype proteomes identified a total of 221 differentially expressed proteins (DEPs) in heterozygotes and 127 DEPs in mutants (Fig. S5A-B; Tables S7,S8). Cyfip2 was the most strongly downregulated protein in mutants, providing a key validation of our unbiased approach (Fig. 3A,S5A; Tables S7,S8). Cyfip2 was slightly but significantly downregulated in heterozygotes as well (Fig. S5B), providing a basis for the sensitization of Cyfip2 heterozygotes to the actin inhibitor CK-869 (Fig. 2B). Cyfip1 expression was not significantly altered in either genotype, indicating that it likely does not act to compensate for the loss of Cyfip2 (Fig. S5A-B). 66 DEPs were shared between *cyfip2* heterozygotes and mutants, while 155 and 61 DEPs were specific to each group, respectively (Fig. S5C). The top 5 upregulated proteins identified in *cyfip2^p400^* heterozygotes in descending order included: microtubule actin cross-linking factor 1 (MACF1), acyl-CoA dehydrogenase family member 11 (ACAD11), cullin 2 (CUL2), calcium channel, voltage dependent, L-type alpha 1S (CACNA1S) and ubiquitin specific peptidase 24 (USP24) (Fig. 3A; top, green). The top 5 downregulated proteins identified in *cyfip2^p400^* heterozygotes in descending order included: tyrosine 3-monooxygenase/tryptophan 5-monooxygenase activation protein theta or 14-3-3 protein theta (YWHAQ), SEC61 translocon subunit alpha 1 (SEC61A1), ribosomal protein S5 (RPS5), enolase 2 (ENO2), and ribosomal protein L18A (RPL18A) (Fig 3A; top, red). The top 5 upregulated proteins identified in *cyfip2^p400^* mutants in descending order included: ACAD11, mitochondrial NADH dehydrogenase 4 (MT-ND4), NIPBL cohesion loading factor b (NIPBL), troponin C2 fast skeletal type (TNNC2), and USP24 (Fig 3A; bottom, green). The top 5 downregulated proteins identified in *cyfip2^p400^*mutants in descending order included: cytoplasmic FMR1-interacting protein 2 (CYFIP2), collagen type VI alpha 3 (COL6A3), stress-induced phosphoprotein 1 (STIP1), thymosin beta (TMSB10/TMSB4X) and collagen type 1 alpha 1 (COL1A1) (Fig 3A; bottom, green). These changes highlight the diverse set of roles that Cyfip2 plays, impacting translational machinery, metabolism, and the extracellular matrix.

**Figure 3.**
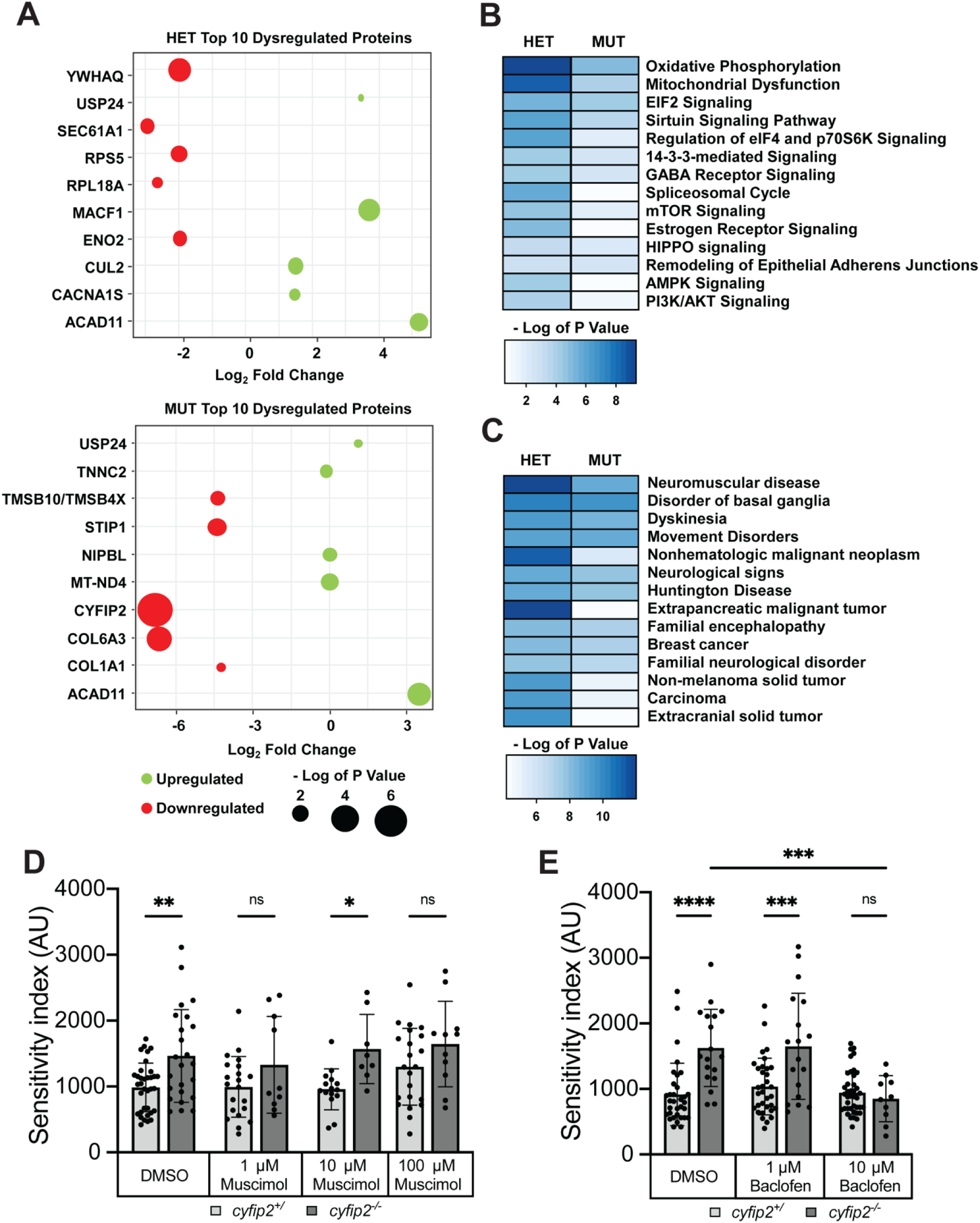
Loss of Cyfip2 causes widespread proteomic changes and GABAB receptor signaling is critical for startle sensitivity. (A) Bubble plots reporting the level of significance of the top 10 dysregulated proteins for both *cyfip2* heterozygous (top) and mutant (bottom) groups compared to wildtype controls. The size of the dot is proportional to the significance of the results while the color code represents the log2 fold change; top five upregulated (green), and top five downregulated (red) proteins are shown. (B-C) Heat maps displaying the impacted canonical pathways (B) and diseases and biological functions (C) from IPA functional analysis. The blue-colored gradient indicates the degree of enrichment for the listed pathways or diseases/functions, as represented by the – log of the P value for each pathway, disease and/or function. (D-E) Sensitivity indices for 5 dpf *cyfip2* sibling (+/) and mutant (−/−) larvae, treated for 60 minutes prior to testing with muscimol (D) or (E) baclofen. All indices (mean ± SD) were compared using a one-way ANOVA with Sidak’s multiple comparisons correction. p* < 0.05, p** < 0.01; p*** < 0.001; p**** < 0.0001.

IPA analysis of DEPs in *cyfip2* heterozygotes and mutants revealed disruption of multiple pathways common to both genotypes. Notable disrupted pathways shared between *cyfip2* heterozygotes and mutants include oxidative phosphorylation, mitochondrial dysfunction, EIF2 signaling, sirtuin signaling, eIF4/p70S6K signaling, 14-3-3-mediated signaling, and GABA receptor signaling (Fig. 3B). Analysis of the most affected diseases and functions for each genotype supports a role for Cyfip2-mediated regulation in neuromuscular disease, disorder of the basal ganglia, dyskinesia, Huntington disease, breast cancer, familial encephalopathy, and neurological disorders (Fig. 3C). These data provide further evidence of the central importance of Cyfip2 for neural function.

Finally, we sought to confirm that these pathways play a functional role in regulating startle sensitivity, and so we focused on GABA receptor signaling as the most likely to contribute to the hyperexcitability of *cyfip2* mutant larvae. We applied agonists of both ionotropic GABA_A_ receptors (muscimol) and metabotropic GABA_B_ receptors (baclofen) for 1 hour prior to startle testing. Muscimol did not have a consistent impact on mutant hypersensitivity at 1, 10, or 100 µM (Fig. 3D), but baclofen induced a clear restoration of normal sensitivity in *cyfip2* mutants at 10 µM (Fig. 3E). Importantly, baclofen application did not cause sedation at this concentration as mutants were equally responsive as untreated siblings, and the same dose of baclofen did not affect startle sensitivity in sibling larvae. Thus, Cyfip2 maintains a normal startle threshold by promoting GABAergic inhibition through the activity of GABA_B_ receptors. Together our findings establish a set of molecular pathways downstream of Cyfip2 that enable the proper processing of acoustic stimuli to limit sensory over-responsiveness.

## Discussion

Through its ability to modulate both the actin cytoskeleton and protein translation, Cyfip2 is well-positioned to be a critical factor for many processes in neurodevelopment [33,34,37–39,45,49]. Further highlighting its importance, Cyfip2 has been implicated in an array of neuropsychiatric and other conditions, including schizophrenia [62], autism [49,63–65], binge eating [66–68], obesity [69], amyotrophic lateral sclerosis (ALS) [70,71], Alzheimer’s disease [72], epilepsy [65,73–76], and cancer [77–82]. Here we focused on Cyfip2’s role in a common endophenotype of schizophrenia and autism, increased acoustic startle responsiveness [4,24,27,28]. By combining conditional transgenesis, CRISPR/Cas9 gene editing, pharmacology, and discovery proteomics in an *in vivo*, vertebrate model system, we found that both actin and translational regulation pathways are required for Cyfip2 to establish and maintain a normal startle threshold. Our data indicate that through these pathways Cyfip2 modulates both excitatory (NMDA receptor) and inhibitory (GABA_B_ receptor) function to establish and maintain proper sensory responsiveness.

### Cyfip2 acts through both Rac1 and FMRP to establish the acoustic startle threshold

The actin regulating function of Cyfip2-Rac1 interactions is essential for many neurodevelopmental processes including neuronal outgrowth and maturation, synapse formation and function, axon guidance and cell migration [38,44,45,83]. The functional importance of Cyfip2-FMRP interactions, though less well characterized than Cyfip1-FMRP interactions, are thought to similarly impact the expression of many key neurodevelopmental proteins that are directly involved in axon growth, synapse maturation, and synaptic plasticity [34,45,84–86]. Here we used a heatshock-inducible expression system to reveal that Cyfip2 requires the ability to interact with both Rac1 and FMRP to establish the acoustic startle threshold early in neurodevelopment (Fig. 1E). The ability of wildtype Cyfip2 to restore normal startle sensitivity in mutant larvae when expression is induced at 30 hpf (Fig. 1D,E), but not at 48, 72, or 96 hpf (Fig. S4) reveals some potential ways that it may affect the underlying neural circuits. Prior to the rescue window, by 8-15 hpf the command-like Mauthner cells have been specified [87] and begun extending their axons (17-18 hpf) and lateral dendrites (22-23 hpf) [88,89]. During the rescue window from ∼30-48 hpf, other neurons in the startle circuit continue to migrate to their final positions in the ventral hindbrain, and the synaptic contacts within the circuit begin to form and mature, including those between the auditory nerve, Mauthner cells, and excitatory Spiral Fiber Neurons (SFNs) [89–91]. Actin dynamics would be required during this time to facilitate neuronal migration, axon and dendrite growth, and synapse formation. These processes would also require precisely regulated RNA translation through complexes like Cyfip2-FMRP-eIF4E to produce the many proteins needed to establish these connections. When we induced Cyfip2 expression at 5 dpf we observed a clear bimodal distribution with some mutants remaining hypersensitive and a second population with normal sensitivity (Fig. 2A). This pattern was not observed when Rac1 or FMRP binding was abolished, suggesting that both pathways are also needed for Cyfip2 to modulate the startle circuit in this acute context, which would likely occur through changes in neuronal and/or synaptic function rather than altered connectivity. That Cyfip2 expression between 48-96 hpf did not restore the startle threshold in mutants could be due to insufficient levels, but together our conditional expression experiments indicate that Cyfip2 is able to most reliably function when the circuit is in a less mature state.

Our findings also show that Cyfip2-Rac1 but not Cyfip2-FMRP binding is required for the performance of the startle response, as kinematic parameters including latency, turn angle, and duration were largely restored to normal by wildtype and Δ*FMRP* versions of Cyfip2 but remained altered in Δ*Rac1*-Cyfip2 expressing *cyfip2* mutants (Fig. S1). These data demonstrate that the actin regulation and translation regulation functions of Cyfip2, while both are required in some contexts, also have some non-overlapping roles. This is consistent with findings in zebrafish larvae showing that Cyfip2’s interaction with FMRP is dispensable but that its interaction with the Wave Regulatory Complex (WRC), which like the Rac1-Cyfip2 interaction regulates actin polymerization (Fig. 1B), is required for retinal ganglion cell (RGC) axons to properly navigate to their targets in the contralateral optic tectum [45]. In the startle context, our data showing that Δ*FMRP*-Cyfip2 drives a stronger rescue for turn angle and duration than for latency suggest that the Cyfip2-FMRP translational regulation pathway contributes more to the initial sensory processing of acoustic stimuli than the regulation of motor output in the spinal cord. The actin regulatory function of Cyfip2, however, appears to be critical for all of Cyfip2’s known roles in the startle circuit.

### Cyfip2-dependent startle threshold regulation requires FMRP but not FXR1/2

Previously, we observed that mutants from the *fmr1^hu2787^*line have normal startle sensitivity, suggesting that FMRP plays no role in regulating the startle threshold [32]. So here we tested whether Cyfip2, which in contrast to the closely related Cyfip1 has the capacity to also bind with the Fragile X-related proteins FXR1 and FXR2 [33], may instead rely on these binding partners to modulate the startle threshold. Like FMRP, both FXR1 and FXR2 regulate RNA translation [92,93] and are expressed in the brain during early vertebrate neurodevelopment, though divergent expression patterns emerge for the FXR1/2 proteins in later development and adulthood in most vertebrates [94–96]. By analyzing FMRP, FXR1, and FXR2 crispants, however, we found that FXR1 and FXR2 are dispensable but that FMRP is required for normal startle sensitivity (Fig. 1F). This data is consistent with the distinct expression patterns of the FMRP, FXR1 and FXR2 proteins, as well as the inability of FXR1/2 to functionally compensate for the loss of FMRP [94–96]. Similarly, in *Drosophila*, which have only one Fragile X protein family member (dFMR1), re-expressing human FMRP (hFMR1), but not human FXR1 or FXR2, in dFMR1 mutants is sufficient to specifically rescue aberrant neuronal phenotypes [97]. As discussed above, the fact that *fmr1* crispants but not *fmr1^hu2787^*mutants show startle hypersensitivity may be due to genomic adaptation in the ENU-induced *fmr1^hu2787^* line [50]. That we observed that *fmr1* crispants show heightened startle sensitivity in the *cyfip2* mutant background compared to *cyfip2* mutants alone (Fig. 1F) further strengthens our conclusion that Cyfip2 and FMRP work cooperatively to regulate the startle threshold.

### Cyfip2-dependent branched actin dynamics are required for maintaining the acoustic startle threshold

Our rescue experiments indicate that Cyfip2’s actin regulatory function through its binding with Rac1 is required during startle circuit development (Fig. 1E) and that this pathway may also facilitate a more acute role for Cyfip2 in maintaining the startle threshold (Fig. 2A). This conclusion is bolstered by our finding that inhibition of Arp2/3 with 20 µM CK-869 for 30 minutes prior to testing uncovers startle hypersensitivity in *cyfip2* heterozygotes but not wildtypes (Fig. 2B). Thus in wildtypes, Cyfip2 must act acutely to facilitate actin polymerization to maintain the startle threshold in the face of this challenge. Arp2/3-mediated F-actin nucleation creates branched actin filaments, while formin-mediated nucleation produces unbranched filaments [56–58,98]. Our data show that only branched actin nucleation is required for acute maintenance of the startle threshold, while both formin activity and cofilin-mediated actin filament severing play no acute role in regulating the startle threshold (Fig. 2C). Formin 2B has been shown with morpholinos to be required developmentally for normal startle responsiveness, however, and it appears to play a role in the growth of Spiral Fiber Neuron (SFN) axons [60]. SFNs are excitatory interneurons that receive input from the contralateral auditory nerve and project their axons across the midline to the contralateral Mauthner cell, providing a key driving force to initiate the startle response [99]. We previously found that SFNs, but not Mauthner cells, have heightened excitability in *cyfip2* mutants [32], making them a likely place for Cyfip2 to regulate the startle threshold. Functioning acutely, Cyfip2 may impact synaptic input onto SFNs, and it is possible that inhibitory and/or excitatory synapses on SFNs may be modulated by Cyfip2 to maintain the startle threshold. Cyfip1 and Cyfip2 are both enriched at excitatory synapses and regulate dendritic complexity and spine maturation in mouse cortical neurons [84,85]. Both Cyfip1 and Cyfip2 are also found at inhibitory postsynaptic sites in mouse hippocampal neurons, and overexpression of either protein disrupts excitatory/inhibitory (E/I) synaptic balance [86]. It is likely that Cyfip2 functions similarly in the zebrafish startle circuit to regulate neuronal excitability, as our data implicate both excitatory (NMDA receptors; Fig. 2D) and inhibitory (GABA receptors; Fig. 3B,E) pathways.

### Cyfip2 may control sensory processing and other disease-related functions by regulating neurotransmitter receptors, mitochondrial function, and/or cytoskeletal remodeling

Our candidate drug screen to identify potential downstream effectors of Cyfip2 in regulating the startle threshold builds on previous work showing that NMDA receptor function is required for normal startle sensitivity (Fig 2D; Table 1) [61]. While our screen confirmed the known roles of Cl^-^ and K^+^ channels, gap junctions, calmodulin, and PDE4 in regulating startle sensitivity, only the NMDA receptor blocker MK-801 produced a *cyfip2* genotype-specific response, indicating that Cyfip2 may impact NMDA receptor expression and/or function in the startle circuit. The mRNAs of three critical NMDA receptor subunit genes – GRIN1, GRIN2A, and GRIN2B – are all targets of FMRP-mediated translational regulation [100], providing a potential mechanism for the enhanced sensitivity of *cyfip2* mutants to the NMDAR inhibitor MK-801. Our unbiased proteomic analysis of *cyfip2* heterozygotes and mutants did not uncover dysregulation in excitatory synaptic pathways compared to wildtypes, although this may be because we analyzed protein lysates from whole larvae and thus may have diluted out any changes in NMDA receptor expression in specific neuronal subpopulations. It could also be the case that Cyfip2 modulates the membrane localization of NMDA receptors through actin-mediated trafficking rather than impacting total expression levels.

Ingenuity Pathway Analysis (IPA) of our proteomic data revealed that inhibitory GABA receptor signaling is significantly disrupted in *cyfip2* heterozygotes and mutants (Fig. 3B). We confirmed that Cyfip2-mediated regulation of GABA receptor function plays a key role in the startle threshold, showing that activation of GABA_B_ but not GABA_A_ receptors is sufficient to rescue the startle hypersensitivity phenotype in *cyfip2* mutants (Fig. 3D,E). Cyfip2 most likely modulates GABA_B_ receptors through translational regulation via FMRP, as both GABA_B_ receptor transcripts in mouse (GABRAB1 and GABRAB2) are targets of FMRP [100]. Further experiments are needed, though, to determine where and how Cyfip2 affects GABA_B_ receptor expression in the startle circuit. Our data are consistent with recent findings using rats in which baclofen-mediated activation of GABA_B_ receptors restored normal auditory processing in *Cntnap2* knockout animals [101]. GABA_B_ receptors function both pre-synaptically to regulate neurotransmitter release and post-synaptically to activate inward-rectifying K^+^ channels that cause hyperpolarization [102,103]. GABA_B_ receptors are also expressed at high levels throughout the auditory system [104], and baclofen treatment has been shown to improve social avoidance in some individuals with autism [105–107]. It is currently unknown whether baclofen affects sensory processing in clinical populations, however. Our data fit with a growing body of evidence that GABA_B_ receptors are important modulators of auditory function with direct clinical applications.

The most significantly disrupted pathways in our proteomic analysis of *cyfip2* heterozygotes and mutants were oxidative phosphorylation and mitochondrial dysfunction (Fig. 3B). It is unclear if these metabolic functions influence the activity of the acoustic startle circuit, although these pathways are essential within neurons for neuronal development and plasticity, cell death, axon extension and branching, and synaptogenesis [108,109], and so they may also contribute to the many disease associations for Cyfip2 listed above. Our data also show that in the absence of Cyfip2, mitochondrial proteins (ACAD11, MT-ND4) increase in abundance, and cytoskeletal (TMSB10/TMSB4X) and extracellular matrix (ECM) proteins (COL6A3, COL1A1) decrease in abundance (Fig 3A; Table S8). TMSB4X and TMSB10 both suppress actin polymerization [110,111], so their downregulation in *cyfip2* mutants may reflect an attempt to compensate for the loss of Cyfip2- and WRC-mediated actin polymerization. ECM collagens like COL6A3 and COL1A1 are important for multiple aspects of neural development including axon guidance [112,113], and so this may reflect another potential mechanism for Cyfip2’s developmental role in regulating the startle circuit. Our analysis of diseases and functions impacted by the loss of Cyfip2 include multiple neurological and neuromuscular conditions (Fig. 3C). These findings reinforce the known associations between *cyfip2* and ALS [70,71] and Alzheimer’s disease [72], further underscoring the importance of Cyfip2 for neural function beyond the startle circuit. Further work on the links between Cyfip2, its molecular effectors, and the development, function, and maintenance of neural circuits will improve our understanding of and ability to treat these varied conditions.

## Materials & Methods

### Zebrafish Husbandry and Maintenance

All animal use and procedures were approved by the North Carolina State University Institutional Animal Care and Use Committee (IACUC). Zebrafish embryos were obtained from the Zebrafish International Resource Center (ZIRC), the University of Pennsylvania, or generated at North Carolina State University and raised in a recirculating housing system. Animals were fed and housed at a 5 zebrafish/L density under a 14h:10h light-dark cycle at 28°C.

Embryos were generated by a male and female pair placed in a mating box (Aquaneering) containing system water and artificial grass. The following morning, 2-3 hours into the light cycle, embryos were collected in petri dishes containing E3 embryo media (5 mM NaCl, 0.17 mM KCl, 0.33 mM CaCl2·2H2O, 0.33 mM MgSO4 in water). Embryos were examined under a brightfield compound microscope for fertilization and proper development and were kept in groups ≤ 65. Embryos were placed in a 29°C incubator on a 14h:10h light-dark cycle. 50% of E3 media changed daily, and any embryos with gross morphological defects were removed and euthanized.

### DNA Extraction & Genotyping

Fin biopsies were obtained from adult fish anesthetized in 0.02% Tricaine (MS-222; Fisher) in system water. Fin clips were taken using a razor blade to remove ∼2-3 mm of tissue from the tail fin and samples were immediately fixed in 100% MeOH. Larval samples were individually fixed in 100% MeOH following behavioral testing. DNA was extracted using the HotShot DNA lysis method which consisted of a tissue lysis with base solution (25mM NaOH, 0.2mM EDTA), sample incubation at 95°C for 30 minutes, and sample neutralization with neutralizing solution (40mM Tris-HCl). *cyfip2^p400^* fish were genotyped using either dCAPS PCR and restriction digest with ApoI-HF [32] or the rhAmp SNP Genotyping System (IDT). rhAmp SNP genotyping was carried out using *cyfip2* locus and allele specific primers (Table S3) targeting the wildtype and *cyfip2^p400^* alleles. Genotyping for *Tg(hsp70:cyfip2-GFP), Tg(hsp70:cyfip2-(C179R)-GFP)* and *Tg(hsp70:cyfip2-(K723E)-GFP)* was accomplished by PCR amplification with primers specific to GFP (Table S4) followed by agarose gel electrophoresis.

### Molecular Cloning

Alternative *cyfip2* rescue constructs (Δ*Rac1*; Δ*FMRP*) were generated from a pENTR *cyfip2*-EGFP plasmid [32] using custom primers and the Q5 Site Directed Mutagenesis Kit (NEB) to induce the desired C179R (Δ*Rac1*) and K723E (Δ*FMRP*) mutations. Mutagenesis was confirmed using restriction digest and Sanger sequencing (Table S3). LR Gateway Cloning (ThermoFisher) was used to insert the altered *cyfip2-*EGFPs into the pDEST I-SceI hsp70 destination vector. Transgenic lines were created by microinjection into 1-cell stage embryos with a transgenesis mix containing phenol red, I-SceI enzyme, and the pDEST I-SceI hsp70:*cyfip2-*(C179R)-EGFP or pDEST I-SceI hsp70:*cyfip2-*(K723E)-EGFP plasmid.

### Inducible Heatshock Rescue & Imaging

Inducible expression of *cyfip2-EGFP,* as well as C179R and K723E variants, was initiated at 30 hpf by placing dechorionated larvae into 96-well plates and incubating at 38°C for 15 or 40 minutes [32]. Following heatshock, larvae were returned to Petri dishes, and given 4 days of recovery at 29°C. GFP fluorescence was confirmed between 4-6 hours post-heatshock for startle experiments using a Nikon SMZ25 stereo microscope with a GFP bandpass filter and Lumen 200 fluorescence illumination system. For day 5 heatshock rescue experiments, larvae were given 4 hours of recovery at 29°C prior to startle sensitivity testing.

For imaging experiments, larvae were treated as above for transgene expression at 30 hpf and at 29°C for 1 hour recovered in petri dishes in groups ≤ 65. After 1 hour of recovery fluorescence was verified, and larvae were visualized using the stereo microscope system described above and larval images were captured at 1-, 3-, 6-, 18-, 24-, 30-, and 42-hours post-heatshock using a Nikon DS-Qi2 monochrome microscope camera. Image analysis was conducted using FIJI (ImageJ) analysis software to manually define ROIs encompassing the entire larval body, excluding the eye and auto fluorescent yolk sac. Fluorescence intensity values reflect the mean gray values recorded for respective ROIs.

### Chemical Exposures

For all exposures, groups of 10-20 larvae (5 dpf) were incubated for specified periods of between 30 minutes and 16 hours within 35 mm Petri dishes in 2 mL of each drug solution. Drug solutions remained on larvae during startle testing for 30 minutes to 1-hour exposures. For 16-hour incubations larvae first received fresh E3 prior to testing. Following incubation, larvae were placed on the 6×6 acrylic testing grid and run through the acoustic startle assay. CK-869, CK-666, MK-801, *N*-phenylanthranilic acid (NPAA), meclofenamic acid (MA), phenoxybenzamine (POBA), etazolate (ETAZ), BMS 204352, muscimol, and baclofen were obtained from Sigma-Aldrich. SMIFH2, IPA-3, GSK429286 and NSC23766 were acquired from Tocris through Fisher Scientific.

### Behavioral Assays

All larvae were tested at 5 days post-fertilization (dpf) unless otherwise stated. On the day of testing, embryos were thoroughly screened for developmental defects, and those with gross morphological defects were removed prior to behavior testing. *cyfip2^p400^* larvae without inflated swim bladders were not discarded, as *cyfip2* mutant larvae fail to inflate their swim bladders [32]. Larvae were adapted to the testing arena lighting and temperature conditions for 30 minutes prior to testing. As previously described, the behavioral testing system consists of a 36-well acrylic grid attached to an acoustic-vibrational shaker (Bruel-Kjaer), a photron mini UX-50 camera, LED lighting, InfraRed illuminator, and an acrylic IR diffuser [5,32,61].

To test the acoustic startle response, 5 dpf larvae were presented with 60 total stimuli with a 20 second interstimulus interval (ISI), with 10 pseudo-randomized trials at each of the following 6 stimulus intensities: 13.6, 25.7, 29.2, 35.5, 39.6 and 53.6 dB. All stimuli were calibrated using a PCB Piezotronics accelerometer (#355B04) and signal conditioner (#482A21), and voltage outputs were converted to dB using the formula dB = 20 log (V/0.775) [32].

### Behavioral Analysis

All responses in the acoustic startle assay were tracked using FLOTE analysis software [5,32,61]. Short latency C-bends (SLCs) were identified by FLOTE using defined kinematic parameters (latency, turn angle, duration, and maximum angular velocity). Startle sensitivity was calculated by measuring the SLC frequency at each of the six stimulus intensities during the 60-stimulus startle assay. Sensitivity indices were defined as the area under the startle frequency vs. stimulus intensity curve calculated using Prism software (GraphPad).

### Larval Sample Preparation for Proteomics

Larvae from incrosses of *cyfip2^+/p400^* carriers were raised as described above. At 3 dpf, DNA was extracted from larvae using the Zebrafish Embryonic Genotyping (ZEG) apparatus (DanioLab). ZEG samples were used for PCR amplification and then submitted for Sanger sequencing to determine the genotype at the *cyfip2^p400^* locus. At 5 dpf larvae were sorted into pools of 30 larvae for each genotype: homozygous wild type, *cyfip2^p400^* heterozygous, and homozygous mutant. Samples were snap frozen in liquid nitrogen and stored at −80°C. They were then resuspended and lysed in 100 µL ammonium bicarbonate (ABC; pH 8) containing 1% sodium deoxycholate (SDC) using a Branson SLPe Sonicator (40:0.15;4C) delivering two 20 second pulses at 20% amplitude intensity separated by 10 seconds between pulses. Lysates were centrifuged at 10,000 rpm for 5 minutes at 4°C, and the supernatants were retained and quantified using Pierce BCA protein quantification (ThermoFisher; Cat #: 23225) and an IMPLEN NP80 nanophotometer. Lysates were submitted the same-day to the Molecular Education, Technology and Research Innovation Center (METRIC) at NC State University for protein digestion and LC-MS.

### Protein Digestion and LC-MS

Each lysed sample was normalized to 200 µg of protein in 200 µL of ABC/SDC solution. Disulfide reduction was conducted by adding 15 µL 50 mM DTT and incubating at 56°C for 30 minutes. 200 µL of 8M urea in 0.1 M Tris buffer (pH 8) was added and samples were transferred to Vivicon 30kD Molecular Weight Cut-off (MWCO) filters. Samples were centrifuged at 12,000 x g for 10 minutes at 21°C. 200 µL of 8 M urea in 0.1 M Tris buffer (pH 8) was added to the top of each filter, as well as 64 µL 55 mM iodoacetamide (IAA) solution and samples were incubated for 1 hour in the dark at room temperature. Samples were centrifuged at 12,000 x g for 20 minutes. 100 µL of 2 M urea, 10 mM CaCl_2_ in 0.1 M Tris buffer (pH 8) was added and samples were centrifuged at 12,000 x g for 20 minutes. The previous step was repeated twice. 100 µL of 0.1 M Tris buffer (pH 7.5) was added and samples were centrifuged at 12,000 x g for 20-45 minutes. This step was repeated twice, with a 1-hour centrifuge period on the final spin. 200 µL of 0.02 µg/mL trypsin in 0.1 M Tris buffer (pH 7.5) was added, and samples incubated overnight at 37°C. Following protein digestion with trypsin, samples were placed into fresh microcentrifuge reservoirs, and 50 µL of quench solution (0.001% zwittergent3-16 in water, 1% formic acid) was added and samples centrifuged at 12,000 x g for 1 hour. 450 µL quench solution was applied to each filter and samples were centrifuged at 14,000 x g for 1 hour. Solutions were dried using a speedvac concentrator and samples were stored dry until LC-MS. Samples were reconstituted in 100 µL of mobile phase A (98% water, 2% acetonitrile, 0.1% formic acid) and peptides were quantified via Pierce BCA assay. Samples were normalized to the lowest peptide concentration for every sample and nanoLC-MS was conducted using a Thermo Orbitrap Exploris 480. This work was performed in part by METRIC at NC State University, which is supported by the State of North Carolina.

### Proteomics data analysis

Shotgun proteomics raw files were processed and quantified with MaxQuant (version 2.2.0.0). Briefly, the built-in Andromeda search engine scored MS2 spectra against fragment masses of tryptic peptides derived from a *Danio rerio* reference proteome containing 93,351 entries including isoforms (UniProt, accessed March 22, 2019). Our database search required variable modifications (methionine oxidation and N-terminal acetylation) and a fixed modification (cysteine carbamido-methylation) along with a minimum peptide length of 7 amino acids and limited the search space to a maximum peptide mass of 4600 Da and a maximum of two missed cleavages. The false discovery rate was controlled with a target-decoy approach at less than 1% for peptide spectrum matches and protein group identifications.

### Bioinformatics

Label-Free quantification (LFQ) intensities from MaxQuant were imported into Perseus software (version 2.0.7.0) and transformed to logarithmic scale with base two. Missing values were replaced with values from the normal distribution, reducing the distributions to a factor of “0.3″ (width) and down-shifting by “1.8″ standard deviations while simulating random values to replace the missing values. This protein quantification was used to measure the fold-enrichment between *cyfip2^p400^* heterozygous/homozygous and *cyfip2* wildtype groups. Statistical significance was calculated using a two-way Student t-test and FPR (p<0.05). Differentially expressed proteins (DEPs) were submitted to ingenuity pathway analysis (IPA) to identify their function, specific processes, and related enriched pathways/diseases.

### Statistical Methods

All statistical analyses were performed using Prism (GraphPad). All data sets were tested for normality in Prism. Subsequent parametric or non-parametric tests and post-hoc analyses were performed using Prism, and significance values (p < 0.05) were reported.

## Acknowledgements

We would like to thank Leonard Collins and Taufika Williams from METRIC at NC State University for their assistance with proteomic analysis. A special thank you to Kimberly Scofield and Kara Carlson for feedback on the manuscript.

## Author contributions

Conceptualization: JCD and KCM; Data curation: JCD, RAM; Formal analysis: JCD, RAM, RGT, SHS, DCC, KCM; Funding acquisition: KCM; Investigation: JCD, RPG, LAR, JAJ, RAM, RGT, SHS, DFB, DCC, KCM; Methodology: JCD, RAM, KCM; Project administration: KCM; Supervision: JCD, DFB, DCC, KCM; Visualization: JCD, RAM, KCM; Writing – original draft: JCD; Writing – review and editing: JCD, KCM

