## Supplementary material for "Cyfip2 controls the acoustic startle threshold through FMRP, actin polymerization, and GABA_B_ receptor function": Fig. s1

**Supporting Information for Deslauriers et al.**

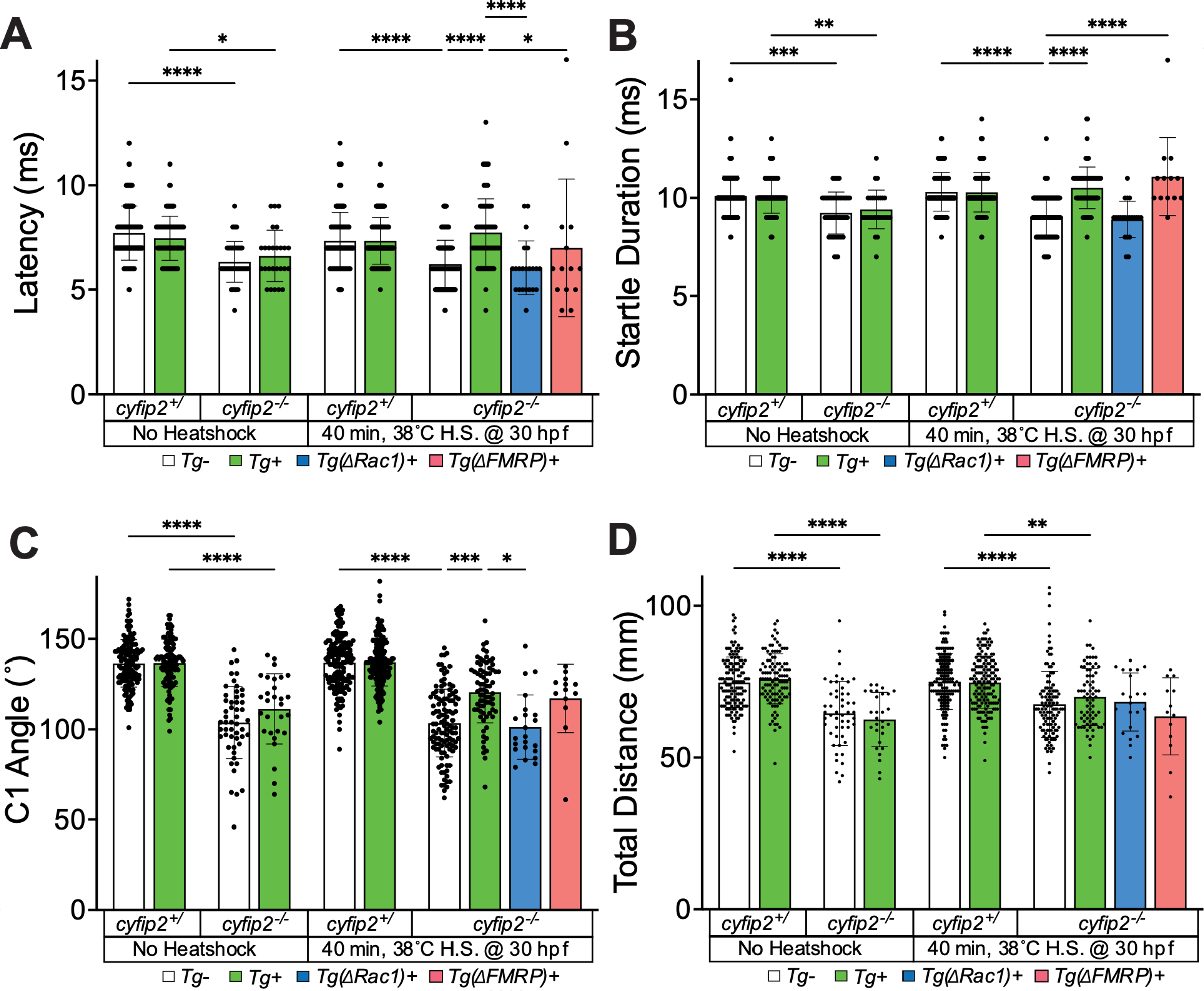

**Figure S1. Restoring Cyfip2 at 30 hpf rescues startle latency, angle, and duration phenotypes in *cyfip2* mutants.**

**(**A) Average startle latency (milliseconds; ms) of larvae shown in Figure 1 at 53.6 dB for *cyfip2* sibling (+/) and mutant (+/+) larvae expressing either normal (*Tg*+; green), Rac1- (∆*Rac1*+; blue) or FMRP/eIF4E- (∆*FMRP*+; pink) binding deficient versions of Cyfip2-EGFP. Comparisons were made to both non-transgenic (*Tg*-) and non-heatshocked controls. All indices (mean ± SD) compared using a Kruskal-Wallis test with Dunn’s multiple comparisons correction; p* < 0.05; p**** < 0.0001. (B) Average startle duration (ms) of larvae shown in Figure 1 at 53.6 dB. Comparisons made as previously. p** < 0.01; p*** < 0.001; p**** < 0.0001. (C) Average initial turn (C1) angle (degrees) of larvae shown in Figure 1 at 53.6 dB. Comparisons made as previously. p* < 0.05; p*** < 0.001; p**** < 0.0001. (D) Total distance traveled (mm) of larvae shown in Figure 1 at 53.6 dB. Comparisons made as previously. p** < 0.01; p**** < 0.0001.

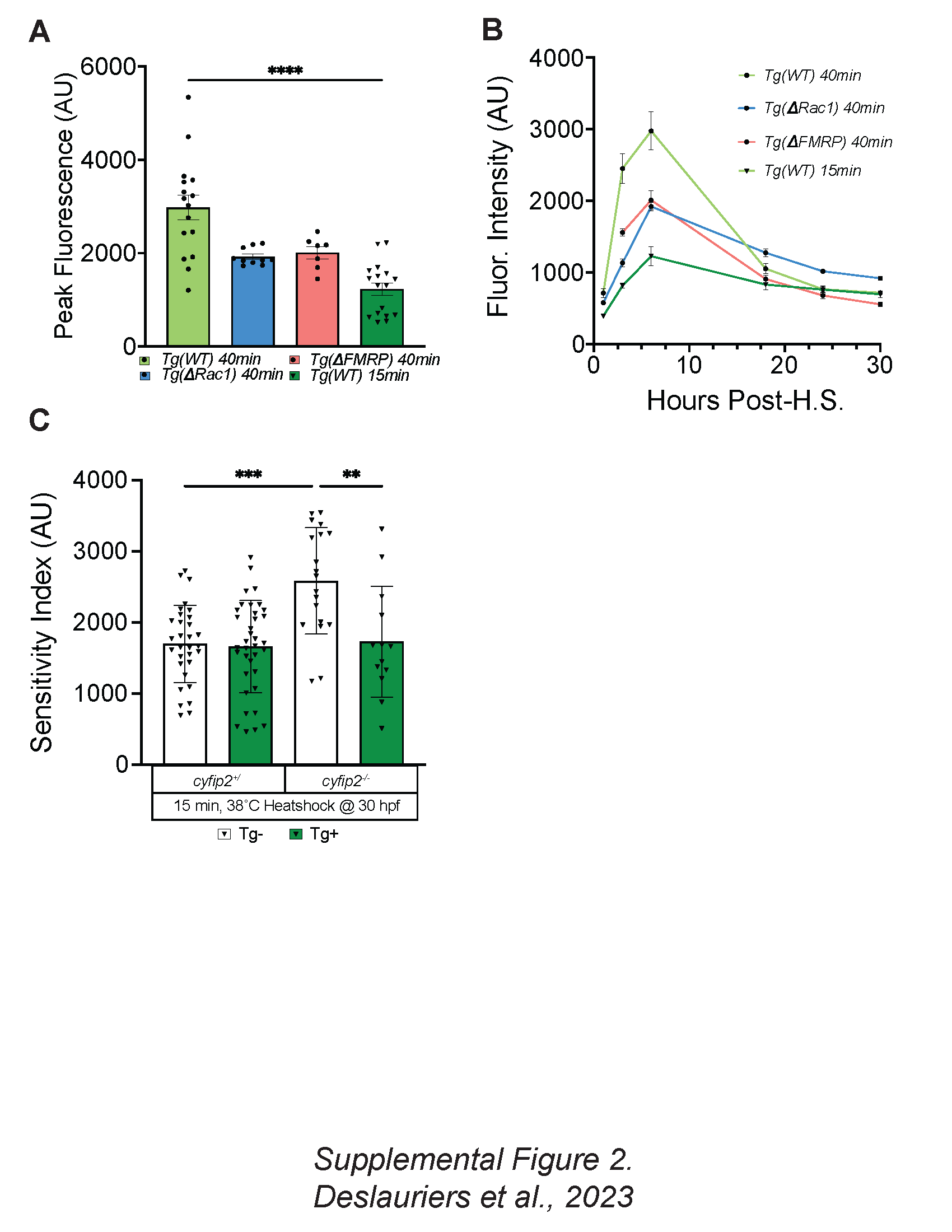

**Figure S2. Rac1 and FMRP/eIF4E interactions are critical to establishing the innate acoustic startle threshold.** (A) Peak fluorescence, at approximately 6 hours post-heatshock of *Tg(hsp70:cyfip2-EGFP)+*, *Tg(hsp70:cyfip2-(∆Rac1)-EGFP)+*, and *Tg(hsp70:cyfip2-(∆FMRP)-EGFP)+* larvae following either a 15- or 40-minute, 38˚C heatshock at 30 hpf. Measurements taken correspond to body specific expression, excluding saturating transgenic expression in the eye and auto fluorescence in the yolk sac, using FIJI image analysis software on lateral view images at the 6-hour post heatshock time point. *Tg*+ 15 min heatshock (green bar, filled triangles), *Tg*+ 40 min heatshock (green bar, filled circles), *Tg*(∆*Rac1*)+ 40 min heatshock (blue bar, filled circles), and Tg(∆*FMRP*)+ 40 min heatshock (pink bar, filled circles). Peak fluorescence values (mean ± SEM) were compared using a Kruskal-Wallis test with Dunn’s multiple comparisons correction; p**** < 0.0001. (B) Fluorescence intensity (mean ± SEM) of *Tg(hsp70:cyfip2-EGFP)+*, *Tg(hsp70:cyfip2-(C179R)-EGFP)+*, and *Tg(hsp70:cyfip2-(K723E)-EGFP)+* larvae following either a 15- or 40-minute, 38˚C heatshock at 30 hpf. Measurements taken as in S2A at 1, 3, 6-, 18-, 24-, and 30-hours post heatshock. Colors as in S2A. (C) Sensitivity indices for 5 dpf *cyfip2* sibling (+/) and mutant (-/-) larvae, following a 15-minute heatshock at 30 hpf to express normal (*Tg*+) Cyfip2-EGFP. Comparisons were made both between transgene conditions and between genotypes. All indices (mean ± SD) compared using a Kruskal-Wallis test with Dunn’s multiple comparisons correction; p** < 0.01; p*** < 0.001.

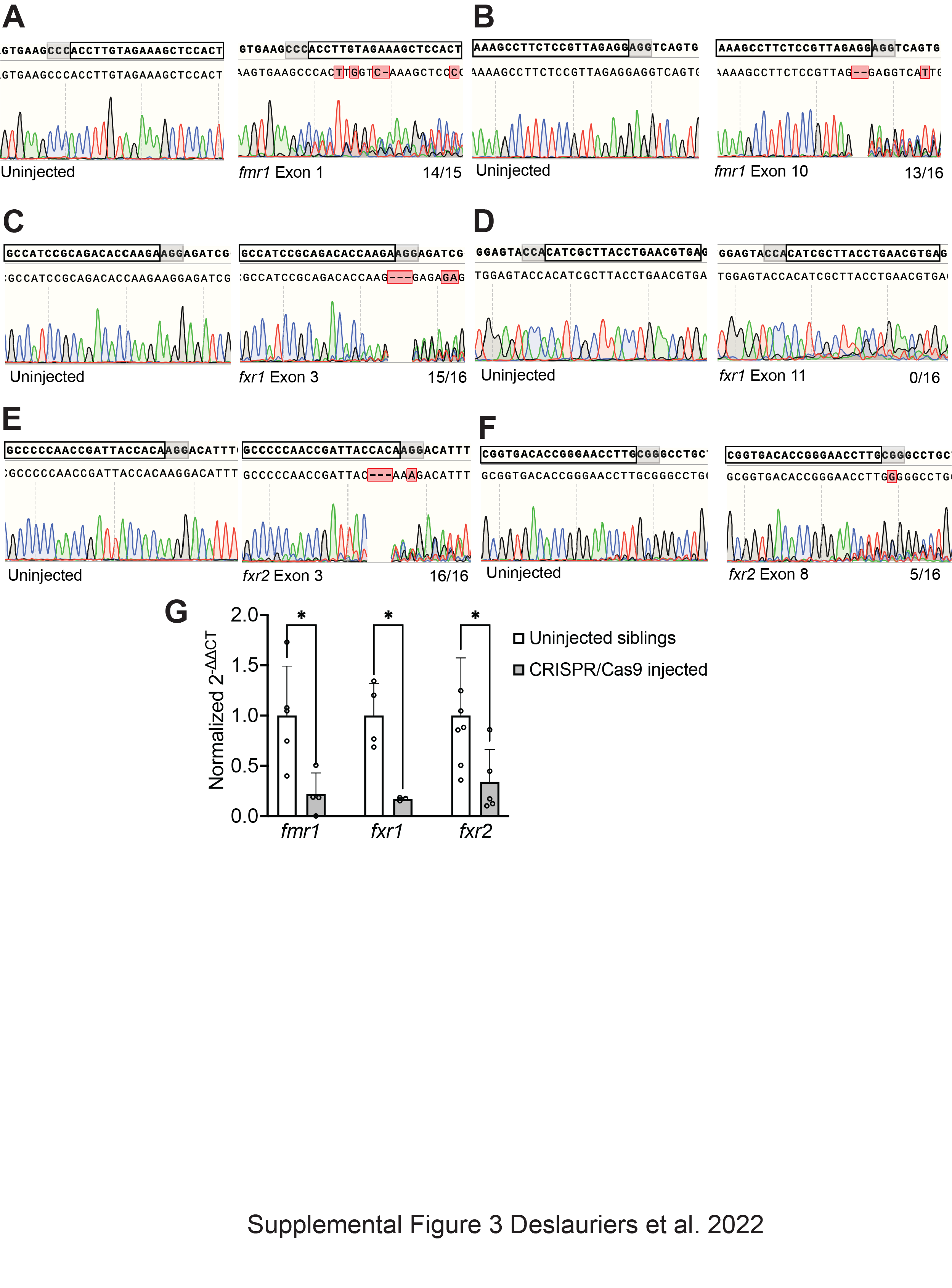

**Figure S3. FMR1, FXR1 and FXR2 CRISPR gRNAs induce mutations at their target sites.** Sanger sequencing chromatogram results for uninjected and CRISPR/Cas9-injected sibling larvae. Target site outlined above each chromatogram by black box, PAM site highlighted in gray, and fractions indicated editing efficiency (# of individuals with edits / total # analyzed). (A) *fmr1* exon 1 target site, (B) *fmr1* exon 10 target site, (C) *fxr1* exon 3 target site, (D) *fxr1* exon 11 target site, (E) *fxr2* exon 3 target site, (F) *fxr2* exon 8 target site. (G) qPCR results showing knockdown of *fmr1, fxr1,* and *fxr2* in CRISPR/Cas9-injected larvae compared to uninjected siblings.

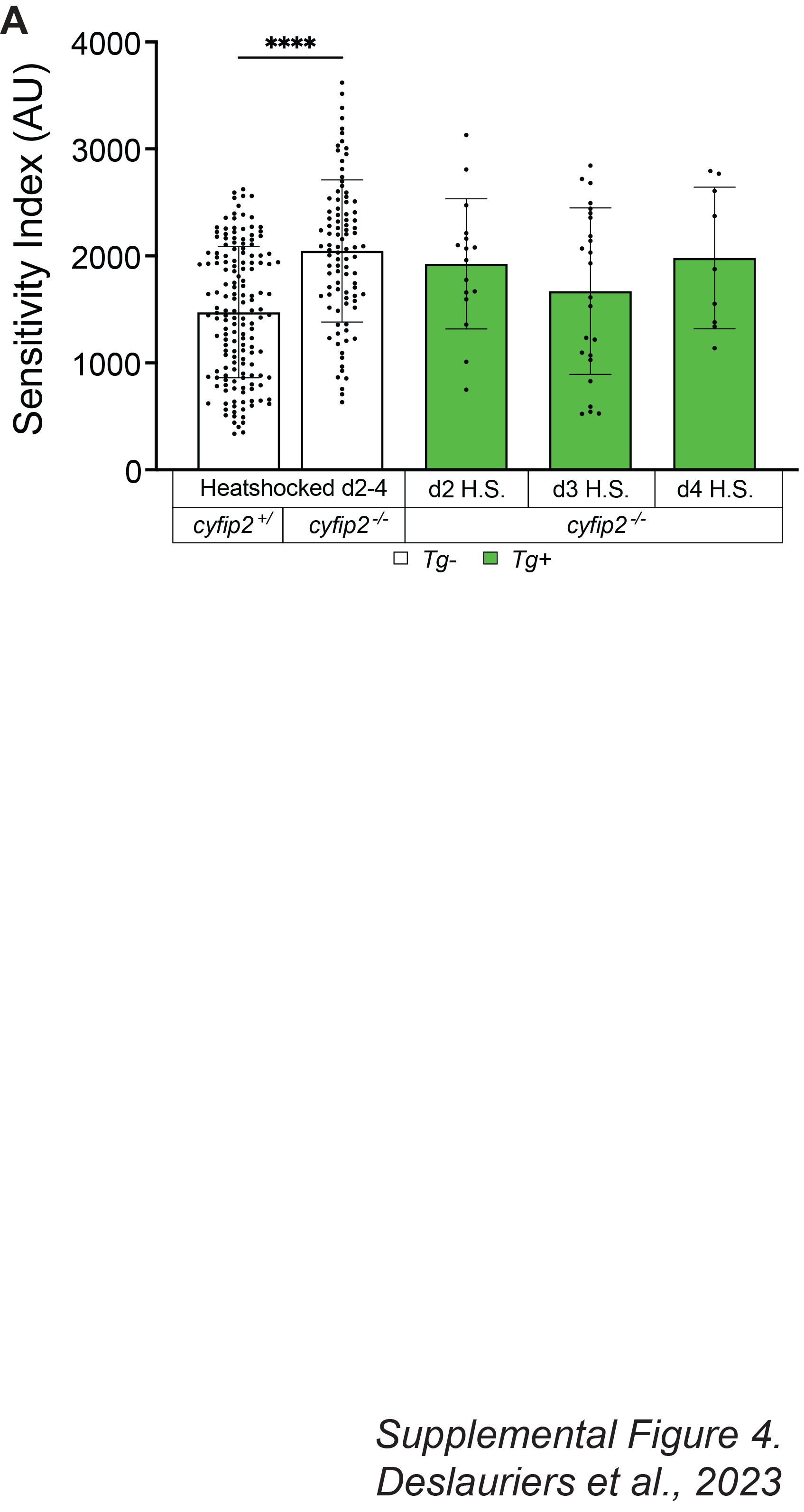

**Figure S4. Cyfip2 fails to rescue mutant hypersensitivity when expressed after 30 hpf. (**A) Sensitivity indices for 5 dpf *cyfip2* sibling (+/) and mutant (-/-) larvae, following a 40-minute, 38°C heatshock at 2, 3 or 4 dpf to express normal (*Tg*+) Cyfip2-EGFP. Comparisons were made both between genotypes, and within genotype by condition. All indices (mean ± SD) compared using a Kruskal-Wallis test with Dunn’s multiple comparisons correction; p**** < 0.0001.

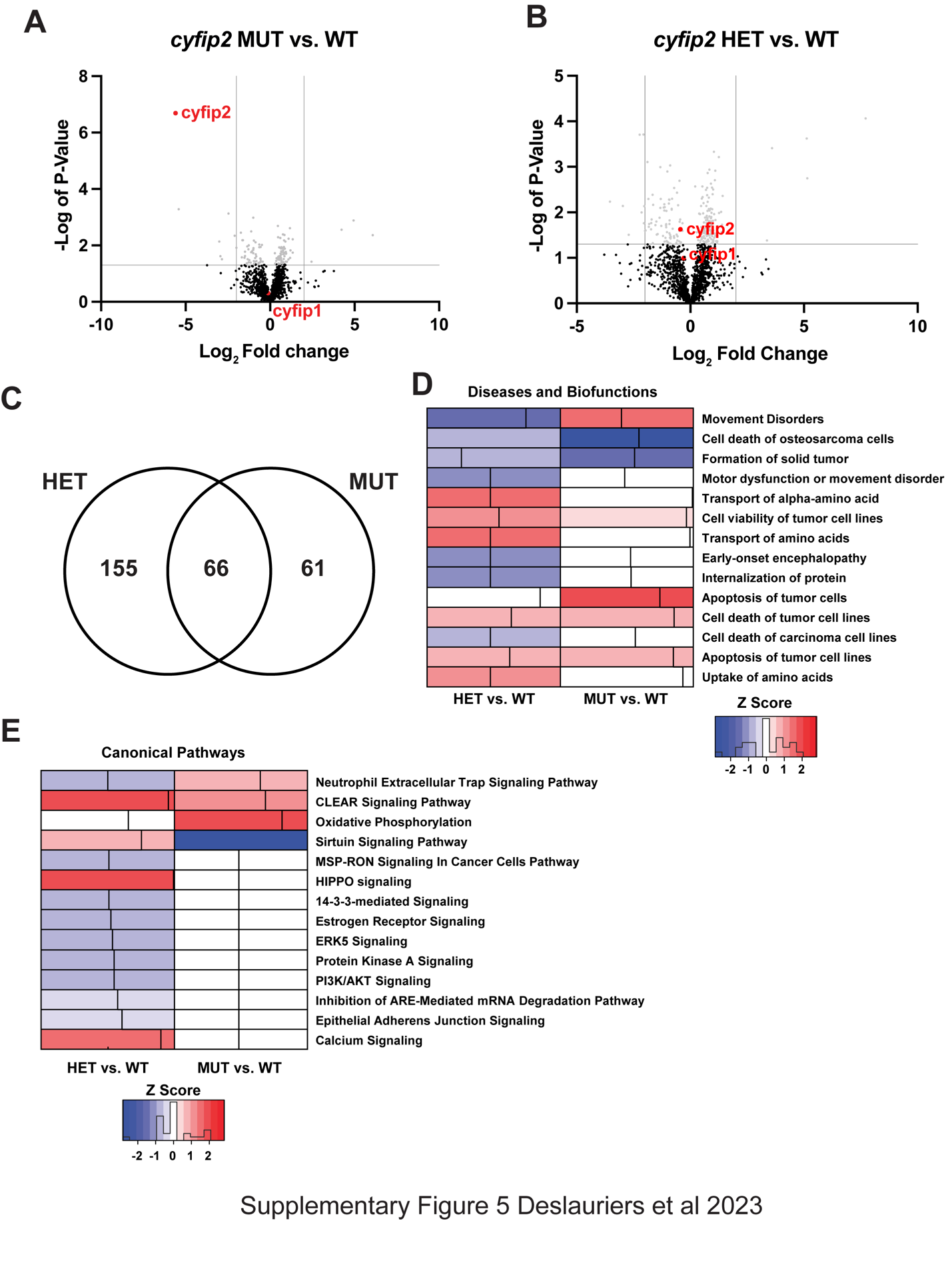

**Figure S5. Loss of Cyfip2 causes substantial changes to the proteome. (**A-B) Volcano plot highlighting the differentially expressed proteins (DEPs) in *cyfip2* mutants (A) and heterozygotes (B) compared to wildtypes. Gray dots indicate the significantly dysregulated proteins (p ≤ 0.05) while the black dots represent non-significantly (p ≥ 0.05) regulated proteins compared to the control group (*cyfip2* wildtype). Red dots highlight Cyfip1 and Cyfip2 proteins. (C) Venn diagram shows the overlap of DEPs between both genotype groups. (D-E) Heat map displaying the impacted diseases and biological functions (D) and canonical pathways (E) identified using IPA to compare *cyfip2* heterozygotes (HET) and mutants (MUT) to wildtypes (WT). The red- or blue-colored rectangles indicate the z-score activities, where red shading indicates predicted activation and blue shading indicates predicted inhibition.

**Table S1. Non-branched actin regulation does not influence acoustic startle regulation.**

| **Compound** | **Concentration (µM)** | **Incubation Period** | | | | |
| --- | --- | --- | --- | --- | --- | --- |
|  |  | **30 min.** | **1 hr.** | **16 hr.** | | |
| SMIFH2 | 1 | 79% of Control, p > 0.99 | 110.13% of Control, p > 0.99 | 97.53% of Control, p > 0.99 | | |
|  | 5 | 69.45% of Control, p = 0.1598 | 109.36% of Control, p > 0.99 | **Lethal** | | |
| IPA-3 | 20 | 75.64% of Control, p = 0.5816 | 134.28% of Control, p = 0.3003 | **Lethal** | | |
|  | 50 | 82.92% of Control, p > 0.99 | 147.04% of Control, p = 0.2631 | **Lethal** | | |
| GSK429286 | 100 | 93.11% of Control, p > 0.99 | - | **62.37% of Control, p*** = 0.0003** | | |
|  |  |  |  | ***cyfip2(+/+)*** | ***cyfip2(+/-)*** | ***cyfip2(-/-)*** |
|  |  |  |  | 62.23% of Control, p = 0.6254 | 78.17% of Control, p > 0.99 | 76.31% of Control, p > 0.99 |

Startle indices comparisons for all tested concentrations and incubation periods of non-branched actin nucleation (SMIFH2) and actin severing pathway antagonists (IPA-3; GSK429286). Significant differences (p < 0.05) between treatment groups and controls are listed (**bold**), using a Kruskal-Wallis test with Dunn’s multiple comparisons correction.

**Table S2. Cyfip2-dependent branched actin polymerization regulates the acoustic startle threshold.**

| **Compound** | **Concentration (µM)** | **Incubation Period (30 minutes)** | |
| --- | --- | --- | --- |
|  |  | *cyfip2^+/+^* | *cyfip2^+/p400^* |
| CK-666 | 5 | 58.42% of DMSO control, p > 0.99 | 110.49% of DMSO control, p > 0.99 |
|  | 10 | 78.97% of DMSO control, p > 0.99 | 120.27% of DMSO control, p > 0.99 |
|  | 20 | 128.33% of DMSO control, p > 0.99 | 154.19% of DMSO control, p = 0.3823 |
|  | 35 | 165.57% of DMSO control, p = 0.7021 | 203.72% of DMSO control, p = 0.1073 |
|  | 50 | **341.10% of DMSO control, p*** = 0.0002** | **421.21% of DMSO control, p**** < 0.0001** |
| CK-869 | 5 | 110.31% of DMSO control, p > 0.99 | 115.61% of DMSO control, p > 0.99 |
|  | 7.5 | 93.81% of DMSO control, p > 0.99 | 84.02% of DMSO control, p > 0.99 |
|  | 10 | 180.76% of DMSO control, p = 0.6087 | **266.15% of DMSO control p*** = 0.0005** |
|  | 20 | 153.06% of DMSO control, p = 0.7128 | **289.66% of DMSO control, p**** < 0.0001** |
|  | 35 | **251.53% of DMSO control, p*** = 0.0006** | **367.73% of DMSO control, p**** < 0.0001** |
|  | 50 | **244.93% of DMSO control, p* = 0.0338** | **339.15% of DMSO control, p**** < 0.0001** |

Startle indices comparisons for all tested concentrations of Arp2/3 inhibitors (CK-666 & CK-869) used in this work. Significant differences (p < 0.05) between treatment groups and controls are listed (**bold**), using a Kruskal-Wallis test with Dunn’s multiple comparisons correction.

**Table S3.** Research animals/strains used in this work.

| **Experimental Models** | **Origin** | **ZFIN ID** |
| --- | --- | --- |
| Zebrafish: *cyfip2^p400^* | Marsden et al., 2018 | ZDB-ALT-180305-9 |
| Zebrafish: *Tg(hsp70:cyfip2-EGFP)* line 14 | Marsden et al., 2018 | ZDB-ALT-180309-1 |
| Zebrafish: *Tg(hsp70:cyfip2-(C179R)-EGFP)* | This Paper | ZDB-ALT-220719-3 |
| Zebrafish: *Tg(hsp70:cyfip2-(K723E)-EGFP)* | This Paper | ZDB-ALT-220719-4 |

**Table S4.** Oligonucleotide Primers used in this work.

| **Primers** | **Sequence (5’ -> 3’)** | **Ref #** |
| --- | --- | --- |
| rhAmp *cyfip2^p400^* Allele Specific #1 | ACCATCTGCTACACAGGTTArUACTG | 192743289; This Paper |
| rhAmp *cyfip2^p400^* Allele Specific #2 | CCATCTGCTCACACAGGTTTrUACTG | 192743288; This Paper |
| rhAmp *cyfip2^p400^* Locus Specific | GCTGCATTTTCATTCCTCTTCTCTCTTTrCTCTC | 192743287; This Paper |
| *cyfip2^p400^* dCAPS Forward | CAAAGTCTTGCTGCGGATAAAAG | Marsden et al., 2018 |
| *cyfip2^p400^* dCAPS Reverse | CTGCACCATCTGCTCACACAAATT | Marsden et al., 2018 |
| *cyfip2^p400^* Reverse | CTCTGAGCGCCAGGTCAAAC | This Paper |
| GFP Forward | GACGTAAACGGCCACAAGTT | Marsden et al., 2018 |
| GFP Reverse | GAACTCCAGCAGGACCATGT | Marsden et al., 2018 |
| *cyfip2* Seq1F | TGGAGGTGATCCCAGGTTAT | This Paper |
| *cyfip2* Seq2F | GGTCTGGACAGTCAGAAGTCTGAT | This Paper |
| *cyfip2* Seq3F | CCCCCTAATGACCCTTGTCTG | This Paper |
| *cyfip2* Seq4F | GCCCTCACAAAATTCAAGAAGCAG | This Paper |
| *cyfip2* Seq5F | GCCAACCACAACGTCTCTGC | This Paper |
| *cyfip2* Seq6F | AGAGACTCGGGACTCCACAG | This Paper |
| *cyfip2*-C179R Fwd | GAACATGAAGcgcAGTGTAAAAAATG | This Paper |
| *cyfip2-*C179R Rev | TTCAGTTCGTCCAGCACA | This Paper |
| *cyfip2*-K723E Fwd | CCTCCTAGACgAACGCTTCCG | This Paper |
| *cyfip2*-K723E Rev | ACACTTCCAGCCATTGCTTT | This Paper |

**Table S5.** List of all CRISPR-Cas9 guide RNAs used in this work.

| **CRISPR gRNA** | **Sequence (5’ -> 3’)** | **Target Exon** | **Ref #** |
| --- | --- | --- | --- |
| *fmr1* gRNA1 | AGTGGAGCTTTCTACAAGGTGGG | Exon 1 | This Paper |
| *fmr1* gRNA2 | AAAGCCTTCTCCGTTAGAGGAGG | Exon 10 | This Paper |
| *fxr1* gRNA1 | GCCATCCGCAGACACCAAGAAGG | Exon 3 | This Paper |
| *fxr1* gRNA2 | TCACGTTCAGGTAAGCGATGTGG | Exon 11 | This Paper |
| *fxr2* gRNA1 | GCCCCCAACCGATTACCACAAGG | Exon 3 | This Paper |
| *fxr2* gRNA2 | CGGTGACACCGGGAACCTTGCGG | Exon 8 | This Paper |

**Table S6.** List of all pharmacologic compounds used in this work.

| **Compound** | **Molecular Target** | **CAS #** | **Manufacturer** |
| --- | --- | --- | --- |
| CK-869 | Arp2/3 antagonist | 388592-44-7 | Sigma-Aldrich |
| CK-666 | Arp2/3 antagonist | 442633-00-3 | Sigma-Aldrich |
| MK-801 | NMDA receptor antagonist | 77086-22-7 | Sigma-Aldrich |
| SMIFH2 | Formin antagonist | 340316-62-3 | Fisher/Tocris |
| IPA-3 | p21-associated kinase 1 (Pak1) antagonist | 42521-82-4 | Fisher/Tocris |
| GSK429286 | Rho associated coiled coil containing kinase (ROCK) antagonist | 37-261-R | Fisher/Tocris |
| *N*-phenylanthranilic acid (NPAA) | Chloride (Cl^-^) channel antagonist | 91-40-7 | Sigma-Aldrich |
| Meclofenamic Acid (MA) | Potassium (K^+^) channel antagonist | 6385-02-0 | Sigma-Aldrich |
| Phenoxybenzamine (POBA) | Alpha-adrenergic receptor/calmodulin antagonist | 62-92-3 | Sigma-Aldrich |
| Etazolate (ETAZ) | Phosphodiesterase 4 (PDE-4) inhibitor | 35838-58-5 | Sigma-Aldrich |
| NSC23766 | Rac1 antagonist | 1177865-17-6 | Fisher/Tocris |
| BMS204352 | K^+^ channel agonist/GABA-A receptor inhibitor | 187523-35-9 | Sigma-Aldrich |
| Muscimol | GABA-A receptor agonist | 2763-96-4 | Sigma-Aldrich |
| Baclofen | GABA-B receptor agonist | 1134-47-0 | Sigma-Aldrich |

**Table S7.** All differentially expressed proteins for the *cyfip2* heterozygous group (HET).

| **HET vs WT** | | |
| --- | --- | --- |
| **Uniprot IDs** | **P value** | **Fold change** |
| A0A2R8RZ61 | 8.66E-05 | 7.71476 |
| Q7ZUM0 | 0.000196 | -2.05927 |
| E7F354 | 0.000197 | -2.2296 |
| B8A4S4 | 0.000238 | 5.11688 |
| E9QFS3 | 0.000389 | 3.59577 |
| B0R1C4 | 0.000465 | 1.03931 |
| F1QT45 | 0.000613 | 1.25343 |
| B7ZD02 | 0.000781 | -1.88612 |
| Q6DGY7 | 0.000926 | -0.88896 |
| A0A0R4IUZ8 | 0.001019 | -1.31951 |
| B8JLV7 | 0.001154 | 0.83548 |
| Q6NUT5 | 0.00122 | 1.09879 |
| Q7ZTS3 | 0.001266 | -1.0092 |
| Q6PD99 | 0.001361 | 0.762697 |
| Q7ZVN9 | 0.001786 | 5.1448 |
| Q6DRC1 | 0.001975 | 1.0806 |
| Q804G7 | 0.002031 | -0.479638 |
| Q7ZUU5 | 0.002345 | 0.905212 |
| Q7ZYX4 | 0.00269 | 0.210278 |
| Q1JPZ7 | 0.002694 | 0.527177 |
| X1WD74 | 0.002859 | 0.957053 |
| Q5TZI1 | 0.002897 | 0.740421 |
| F8W4Q9 | 0.002915 | -0.715132 |
| B0S5S9 | 0.003161 | 1.00585 |
| Q6DBT9 | 0.003202 | 0.686377 |
| Q6PC82 | 0.003687 | -0.591262 |
| Q7SYE1 | 0.003993 | 0.981394 |
| Q7SZC9 | 0.004124 | -1.66381 |
| Q6PBQ4 | 0.004184 | 0.73479 |
| E9QCA9 | 0.004355 | 1.42082 |
| Q804C3 | 0.004622 | 0.54113 |
| F1QUW4 | 0.004741 | 0.814594 |
| A0A2R8QJJ5 | 0.005004 | 0.500837 |
| A8WG05 | 0.005058 | -0.800756 |
| B3DJH0 | 0.005072 | -1.30422 |
| Q6IQQ0 | 0.005082 | -1.84776 |
| Q6DGY4 | 0.005253 | -0.93853 |
| Q6AXJ2 | 0.005264 | -0.920406 |
| Q7SXQ3 | 0.00584 | -3.53452 |
| Q90ZM2 | 0.005852 | 0.760472 |
| B8JL35 | 0.006264 | 0.576817 |
| Q6P027 | 0.006418 | -0.948851 |
| Q7ZV04 | 0.006652 | -1.14902 |
| Q6DG71 | 0.006826 | -1.63392 |
| B0S754 | 0.006975 | -0.540343 |
| Q90ZM2 | 0.007239 | -2.96893 |
| Q803J3 | 0.007245 | 0.474805 |
| E9QE47 | 0.007272 | 0.700467 |
| Q801M0 | 0.0073 | -2.03043 |
| B2GSH6 | 0.007691 | -1.12915 |
| Q6P2T0 | 0.007844 | -1.1738 |
| Q6IQ97 | 0.007976 | -1.31667 |
| F1QMZ6 | 0.00826 | 0.892981 |
| Q6PBX8 | 0.008349 | -0.822399 |
| A5WV37 | 0.008488 | 0.811984 |
| Q1LYP4 | 0.008545 | 0.962926 |
| Q6P959 | 0.008691 | -0.820321 |
| Q6PC34 | 0.009116 | 0.845695 |
| Q90WX5 | 0.009118 | -1.14349 |
| Q6TGY8 | 0.009261 | 0.751429 |
| Q9I9E5 | 0.009283 | -1.0516 |
| F1RAV6 | 0.009806 | 1.35714 |
| P17561 | 0.010064 | -2.14753 |
| A0A8M9P7D1 | 0.010204 | 0.927811 |
| Q6PYX3 | 0.010331 | 1.26646 |
| Q567V5 | 0.010536 | 0.564191 |
| Q7SZD2 | 0.010652 | 0.391492 |
| Q4G5V7 | 0.010925 | 1.0493 |
| D7RVS0 | 0.010925 | 0.429207 |
| B0S553 | 0.011947 | 0.717731 |
| Q6P0H1 | 0.012309 | 0.834073 |
| Q5XJT7 | 0.012654 | 0.921172 |
| F6NK19 | 0.012961 | 0.682257 |
| G4XPM3 | 0.013015 | 0.648204 |
| Q5U3R4 | 0.013168 | -1.35827 |
| Q6NWC6 | 0.01337 | 0.753258 |
| Q6DBW5 | 0.013479 | 0.904397 |
| Q7SY50 | 0.014323 | -1.70132 |
| A0A8M1P182 | 0.01433 | -0.44557 |
| Q5SNP7 | 0.014698 | 1.10813 |
| Q6NWH2 | 0.01483 | 0.689758 |
| F1Q9I7 | 0.014938 | 0.752187 |
| B8A518 | 0.015037 | -0.91227 |
| Q5U369 | 0.015386 | 0.996881 |
| Q6PHE5 | 0.01539 | 0.997368 |
| Q503N6 | 0.015597 | 0.834698 |
| F1QCR7 | 0.015701 | 0.913886 |
| E7FC95 | 0.015893 | -1.49379 |
| Q502L5 | 0.015932 | 0.819911 |
| Q6DGN1 | 0.016077 | -0.470161 |
| A0A2R8RKK2 | 0.016391 | 0.913101 |
| F1QDE7 | 0.016648 | 0.596429 |
| Q803G7 | 0.016773 | 0.605661 |
| Q5XJP7 | 0.01719 | 0.709815 |
| F1Q5B8 | 0.017246 | -1.72005 |
| Q8JHH3 | 0.017291 | -2.04476 |
| O42248 | 0.017431 | -0.343451 |
| F8W4X0 | 0.01804 | -1.81403 |
| A0A8M1NE64 | 0.018124 | -0.923081 |
| Q66L49 | 0.018135 | 0.948545 |
| Q6NWK7 | 0.018146 | -1.06605 |
| Q7ZUD6 | 0.018215 | 0.835702 |
| Q6RKB0 | 0.018704 | 1.39215 |
| Q5U3E5 | 0.019095 | 0.912415 |
| Q8JH72 | 0.019139 | -1.06957 |
| Q4VBK0 | 0.0192 | -1.53365 |
| Q7T341 | 0.019305 | 0.711135 |
| F1R3D3 | 0.019784 | -1.28123 |
| B2GS26 | 0.019955 | 0.786676 |
| A4FUK1 | 0.020276 | 1.20295 |
| Q6PFU0 | 0.020812 | -1.82557 |
| Q1LVM0 | 0.021127 | 0.479118 |
| F8W4E2 | 0.021296 | 1.22339 |
| Q6DGT8 | 0.021298 | -1.0666 |
| Q7T296 | 0.021559 | 0.559891 |
| Q8UVZ4 | 0.021686 | 0.706464 |
| CON__P35527 | 0.021755 | 0.819402 |
| Q6DGS3 | 0.02185 | 0.552607 |
| Q9I8U8 | 0.022048 | 1.22729 |
| Q803L1 | 0.02245 | 0.679484 |
| Q5D018 | 0.022475 | 0.341981 |
| F1QII8 | 0.022878 | 0.595292 |
| Q566S4 | 0.022946 | -0.781161 |
| A7E2L8 | 0.023575 | -1.72753 |
| A5A5E1 | 0.023603 | -0.437627 |
| E7F5G8 | 0.02374 | 0.85531 |
| E7FGW2 | 0.023919 | 0.839511 |
| X1WET9 | 0.024128 | 0.60789 |
| E7FBU7 | 0.024738 | 0.809415 |
| Q6IQK3 | 0.024859 | -0.379822 |
| F6PBX0 | 0.024882 | 1.01335 |
| Q66IB3 | 0.024931 | -1.97329 |
| Q08B95 | 0.025284 | 0.607609 |
| Q803H5 | 0.025368 | -0.478846 |
| Q6P3J5 | 0.02548 | -0.342405 |
| Q6NWI7 | 0.025516 | -1.99737 |
| F1QKW3 | 0.025541 | 0.888355 |
| B0R193 | 0.025913 | 0.675828 |
| Q6AXJ2 | 0.026561 | 0.483077 |
| E7FFL3 | 0.026751 | 0.683697 |
| Q6PBM9 | 0.026821 | 0.365942 |
| F5HSE3 | 0.026872 | 1.37693 |
| Q6PHJ1 | 0.027168 | -1.02579 |
| A5HLY6 | 0.027238 | 0.537177 |
| Q6P0H6 | 0.027449 | 0.285213 |
| Q0P408 | 0.028206 | 0.632262 |
| A4FVL3 | 0.028395 | 0.572223 |
| Q32LR2 | 0.02873 | 0.911809 |
| Q6TNV0 | 0.028779 | 0.63668 |
| A7MBY4 | 0.028987 | -0.920881 |
| U3JAA1 | 0.029106 | 0.78484 |
| F1QPP6 | 0.029141 | 1.07232 |
| Q5TZ35 | 0.029741 | 0.74973 |
| E7EZE6 | 0.030136 | -1.43541 |
| A0A0R4IP12 | 0.030347 | 0.479474 |
| A2CEA9 | 0.030526 | 0.936813 |
| Q6PBJ9 | 0.030622 | 0.893379 |
| F1QTW8 | 0.030716 | 0.701908 |
| Q6Q420 | 0.030796 | 0.848694 |
| Q9DGR5 | 0.030806 | -1.48242 |
| Q7T356 | 0.03086 | -0.602564 |
| Q7ZWJ4 | 0.031077 | -2.71144 |
| Q7SX97 | 0.031287 | 0.821599 |
| Q08BY9 | 0.031438 | 0.713483 |
| Q6DC80 | 0.031539 | -0.88979 |
| A4QN66 | 0.031829 | 0.81361 |
| Q568L3 | 0.03283 | 1.09307 |
| E7F8M1 | 0.032937 | -1.20617 |
| Q7ZUT3 | 0.032972 | 0.722386 |
| Q0R680 | 0.033196 | -0.724326 |
| Q5U3G0 | 0.033329 | 0.96201 |
| Q6DBW7 | 0.033451 | 0.598706 |
| Q802X5 | 0.03365 | -0.566586 |
| Q5RHR9 | 0.034789 | -0.861663 |
| F6P9B5 | 0.035034 | 0.506165 |
| A5A4L9 | 0.035657 | 0.528285 |
| Q503D5 | 0.03579 | 0.861811 |
| Q7T2P3 | 0.035996 | -0.899419 |
| Q1LVE8 | 0.036044 | 1.05433 |
| A2RUZ3 | 0.036188 | 0.565196 |
| F8W2Z3 | 0.036363 | -0.709115 |
| A7E2K5 | 0.036366 | -1.0977 |
| F1Q8W8 | 0.037522 | 0.642242 |
| C5IG48 | 0.037925 | -0.867814 |
| Q6DGE9 | 0.038805 | 0.494877 |
| Q8UUX9 | 0.038886 | 0.923285 |
| O42271 | 0.03978 | 0.776283 |
| Q1LVQ8 | 0.040416 | 0.993941 |
| Q6IQI2 | 0.040487 | 1.01537 |
| Q6P6E0 | 0.040686 | 0.347212 |
| Q5XIZ4 | 0.04114 | 0.773116 |
| Q6NWJ2 | 0.041379 | -0.573219 |
| A8E7T1 | 0.041641 | 3.37243 |
| Q6PBW7 | 0.042433 | 0.446904 |
| X1WCE0 | 0.042587 | 1.13429 |
| F1QKX8 | 0.042624 | 0.709089 |
| E9QG51 | 0.042638 | -1.04685 |
| Q5U3U1 | 0.042803 | 1.00916 |
| A0A8M9PPE1 | 0.043044 | 0.56731 |
| B2GPU7 | 0.043045 | -1.04725 |
| Q5U396 | 0.043053 | 1.18915 |
| E9QFU8 | 0.043186 | 0.804141 |
| Q1LWH1 | 0.043812 | -0.489061 |
| Q4G5T8 | 0.043931 | 1.14432 |
| Q9IAB6 | 0.043991 | 0.986611 |
| Q9MIY5 | 0.044195 | -1.13639 |
| B5DDZ4 | 0.044323 | 0.736048 |
| D5LHQ7 | 0.044385 | 0.609837 |
| Q68EH2 | 0.044392 | -1.00263 |
| O57521 | 0.045903 | -0.411109 |
| CON__P00761 | 0.046359 | -0.968283 |
| B2GSX0 | 0.046508 | 0.669394 |
| Q7T3G2 | 0.046605 | -0.609184 |
| Q5RLN6 | 0.047349 | 0.425562 |
| Q6DH38 | 0.047581 | -1.6536 |
| Q567D7 | 0.047841 | 1.00903 |
| Q6TLF6 | 0.047968 | 1.17672 |
| Q4G5K8 | 0.048197 | 0.927052 |
| I3ITF4 | 0.048443 | -0.886983 |
| D2X2I2 | 0.048705 | -0.864377 |
| Q6IQ59 | 0.048911 | -0.431266 |

**Table S8.** All differentially expressed proteins for the homozygous *cyfip2* mutant group (MUT).

| **MUT vs. WT** | | |
| --- | --- | --- |
| **Uniprot IDs** | **P value** | **Fold change** |
| A5A5E1 | 2.03911E-07 | -5.59564 |
| F1QMR9 | 0.00052236 | -5.40953 |
| Q90ZM2 | 0.000742062 | -2.45573 |
| Q7ZTS3 | 0.001033618 | -0.989005 |
| Q7ZVN9 | 0.001309001 | 4.94043 |
| Q15I86 | 0.00205608 | 0.460352 |
| F8W4E2 | 0.002459291 | 1.17022 |
| B8A4S4 | 0.002777218 | 4.23144 |
| Q6IQQ0 | 0.003172488 | -1.1173 |
| Q7ZWJ4 | 0.003522735 | -1.72366 |
| Q6PBQ4 | 0.004052005 | 0.758154 |
| D7RVS0 | 0.004280456 | 0.851676 |
| A0A2R8RZ61 | 0.004334111 | 6.07489 |
| Q8JGR4 | 0.00456983 | -2.09505 |
| A2BE76 | 0.00513949 | 0.47299 |
| Q804C3 | 0.005435131 | 0.606574 |
| F1QT45 | 0.005818754 | 1.08996 |
| Q9MIY1 | 0.006076171 | 1.33827 |
| Q6PFU0 | 0.006509086 | -1.64371 |
| Q6NUW5 | 0.006565231 | -1.61033 |
| Q4G5V7 | 0.006652732 | 0.824453 |
| Q08C47 | 0.006788597 | -0.963854 |
| A5WV37 | 0.0072716 | 0.904072 |
| Q5RKM3 | 0.007343617 | -3.00481 |
| Q6NWK7 | 0.007706373 | -0.855665 |
| Q7ZUP6 | 0.009266378 | -0.639457 |
| B3DJH0 | 0.009579658 | -0.482817 |
| Q6DG71 | 0.009897374 | -1.26208 |
| B0UYS0 | 0.010827046 | -1.35404 |
| Q1L8Q3 | 0.011347495 | 0.816093 |
| B2GRG9 | 0.011405649 | -0.558996 |
| A0N0A7 | 0.01157071 | 0.669225 |
| Q804G9 | 0.011589908 | -0.64098 |
| Q9YH92 | 0.011746812 | 0.542367 |
| Q6PBW7 | 0.011936033 | 0.576003 |
| Q5RZ65 | 0.01204814 | -1.99309 |
| Q66HZ3 | 0.012094001 | 0.78498 |
| F1QMZ6 | 0.012444286 | 0.741557 |
| P87360 | 0.012459482 | -0.987051 |
| Q502L5 | 0.012858491 | 0.655478 |
| E7FC33 | 0.013080972 | 0.876769 |
| B7ZD02 | 0.013674139 | -1.28274 |
| Q7ZUI4 | 0.014325177 | -0.989663 |
| A7E2K4 | 0.014937228 | 0.768817 |
| E9QBF1 | 0.015047349 | 1.20558 |
| Q7SXX4 | 0.016119442 | 0.659585 |
| B2GSH6 | 0.016226319 | -1.07757 |
| Q9DEU1 | 0.016772199 | -1.48524 |
| Q8UUX9 | 0.016879128 | 0.927227 |
| B8JL35 | 0.017280647 | 0.691652 |
| A7MCQ5 | 0.017463448 | -1.11373 |
| F5HSE3 | 0.017940725 | 1.34055 |
| A0A0K0F5S9 | 0.017980013 | 0.809795 |
| E9QCD1 | 0.018926487 | 0.28596 |
| Q6NX91 | 0.018962692 | -0.991968 |
| B0R1C4 | 0.019269924 | 0.955031 |
| B2GS26 | 0.019459425 | 0.84403 |
| Q7ZUY9 | 0.019565908 | 0.621363 |
| F1QJU0 | 0.01992783 | 0.94014 |
| U3JAA1 | 0.020536196 | 0.78544 |
| Q8JHG2 | 0.020872766 | -0.845503 |
| Q567V5 | 0.020991332 | 0.653493 |
| A7E2L8 | 0.023647215 | -1.48568 |
| Q803G7 | 0.023807292 | 0.605124 |
| Q45QT2 | 0.024222 | -2.9438 |
| Q803G1 | 0.024300209 | 0.534508 |
| A4QN66 | 0.024331563 | 0.850256 |
| Q7T1R2 | 0.024443872 | -0.816371 |
| Q6NSN5 | 0.025136222 | 0.485523 |
| Q6TGT0 | 0.026478904 | -1.66869 |
| Q6PC34 | 0.026535666 | 0.61495 |
| Q6DRD6 | 0.027525169 | 0.55623 |
| Q6P271 | 0.027539116 | -1.51148 |
| A5A4L9 | 0.027549264 | 0.453133 |
| Q6RKB0 | 0.027945379 | 1.17807 |
| Q6IMF9 | 0.027991747 | -0.880959 |
| Q7SX97 | 0.028174745 | 0.651189 |
| Q7ZSY3 | 0.028547626 | 0.383965 |
| F1QXB0 | 0.02909913 | -0.58536 |
| A0A8M1NUM0 | 0.030134223 | 0.536331 |
| E7FGU0 | 0.030396949 | -0.564946 |
| E7FBD3 | 0.030633033 | 0.810468 |
| Q6PHJ4 | 0.030688807 | -1.32292 |
| F6NK19 | 0.030819103 | 0.465742 |
| B1WB89 | 0.030951379 | -2.85524 |
| Q5RGQ4 | 0.031226981 | 0.564689 |
| Q7SYI7 | 0.031303291 | 1.08607 |
| Q6P6E0 | 0.032237399 | 0.543235 |
| F1R922 | 0.033585359 | -1.23018 |
| Q7T160 | 0.033845427 | 0.674531 |
| B8JLR6 | 0.035505039 | 0.360222 |
| Q05AP7 | 0.035874842 | -0.8066 |
| Q803G3 | 0.036131833 | -0.96478 |
| Q7ZUP2 | 0.03625434 | -0.551839 |
| A0A0R4ID71 | 0.037442083 | -0.859924 |
| Q08BX5 | 0.037567301 | 0.932148 |
| Q5XJT7 | 0.037568166 | 0.993062 |
| A8E5J3 | 0.037577683 | 0.673695 |
| Q803M8 | 0.037853845 | -0.437536 |
| F1Q6K1 | 0.037923639 | 2.44782 |
| E7F1G8 | 0.039049008 | 0.775294 |
| Q49HM7 | 0.039433924 | -0.583182 |
| Q4VBK0 | 0.039829972 | -1.27735 |
| E9QF63 | 0.041049701 | -1.15528 |
| Q6Q420 | 0.041214497 | 0.796659 |
| B0S5T1 | 0.04128003 | -0.647926 |
| Q08B95 | 0.042082352 | 0.793835 |
| Q6PC82 | 0.042264915 | -0.66847 |
| B8JLV7 | 0.042667775 | 0.69309 |
| Q6TNV0 | 0.043011046 | 0.662924 |
| D2K290 | 0.043024913 | -1.38382 |
| Q1JPZ7 | 0.043664652 | 0.308879 |
| Q6DBW7 | 0.044002768 | 0.492137 |
| Q7ZVF9 | 0.044286361 | -0.697097 |
| Q6DGE9 | 0.044472342 | 0.485659 |
| Q6NYB0 | 0.044772361 | -0.948409 |
| X1WET9 | 0.04550614 | 0.586884 |
| R4GDP5 | 0.045929318 | 0.658113 |
| E7FBU7 | 0.046439764 | 0.566435 |
| Q4V9H6 | 0.046743368 | 0.831491 |
| Q6DGN1 | 0.046922377 | 0.371283 |
| A2VD35 | 0.047314036 | 0.849927 |
| Q7ZVQ3 | 0.047615681 | -0.848595 |
| Q7SXN2 | 0.049150733 | -0.857413 |
| Q9DDE1 | 0.049492569 | 0.910999 |
| F1QUY7 | 0.049811544 | -0.336958 |
| Q6YBS2 | 0.049935569 | -0.368233 |

**Table S9.** List of canonical pathways activation/inhibition for each genotype and their respective z scores.

| **Canonical Pathways** | **HET vs WT** | **MUT vs WT** |
| --- | --- | --- |
| Neutrophil Extracellular Trap Signaling Pathway | -2.333 | 0.816 |
| CLEAR Signaling Pathway | 1.89 | 1 |
| Oxidative Phosphorylation | -0.905 | 1.633 |
| Sirtuin Signaling Pathway | 0 | -2.449 |
| MSP-RON Signaling In Cancer Cells Pathway | -2.236 | N/A |
| HIPPO signaling | 2.236 | N/A |
| 14-3-3-mediated Signaling | -2.236 | N/A |
| Estrogen Receptor Signaling | -2.121 | N/A |
| ERK5 Signaling | -2 | N/A |
| Protein Kinase A Signaling | -1.89 | 0 |
| PI3K/AKT Signaling | -1.89 | N/A |
| Inhibition of ARE-Mediated mRNA Degradation Pathway | -1.633 | N/A |
| Epithelial Adherens Junction Signaling | -1.342 | N/A |
| Calcium Signaling | 1.342 | N/A |
| AMPK Signaling | 1.342 | N/A |
| Endocannabinoid Neuronal Synapse Pathway | -1 | N/A |
| SNARE Signaling Pathway | 1 | N/A |
| Insulin Secretion Signaling Pathway | 0 | 1 |
| Cardiac Hypertrophy Signaling | -1 | N/A |
| Xenobiotic Metabolism CAR Signaling Pathway | -1 | N/A |
| EIF2 Signaling | N/A | -1 |
| G Beta Gamma Signaling | -1 | N/A |
| Spliceosomal Cycle | 0.816 | N/A |
| Coronavirus Pathogenesis Pathway | -0.447 | N/A |
